# Oxidant-mediated activation and inhibition of LRRC8C and LRRC8D channel currents require N-terminal and Transmembrane 1 pore domains

**DOI:** 10.64898/2026.01.22.701128

**Authors:** Jeffrey Rohrbough, Hong N. Nguyen, Fred S. Lamb

## Abstract

Leucine Rich Repeat Containing 8C (LRRC8C) anion channels modulate NADPH oxidase 1 activity and allow extracellular superoxide influx promoting inflammatory signaling. Here we studied chimeric 8C/8D channels and identified oxidant-dependent current modulation within the N-terminus (NT) and first transmembrane domain (TM1). Chloramine-T (ChlT) elicited inhibitory and activating current responses, whereas other redox agents had comparatively little impact. ChlT moderately inhibited wild-type (WT) 8C current and abrogated block by DCPIB. Substitution of the 8D NT (8D_1-22_) conferred ChlT-dependent current activation, as did 8D_2-4_, 8D_5-11_, I2F, and I2Y substitution. *M48T* (distal TM1) substitution enhanced WT 8C current inhibition and impaired activation in NT mutants. An *M48D* mutation diminished 8C current block by DCPIB by ∼50%. WT 8D currents were potently inhibited by ChlT. Substitution of the 8C first extracellular loop (EL1) weakened inhibition, while 8C EL1 + TM1_45-49_ substitution produced ChlT-mediated current activation. 8C_45-49_ or *T48M* substitutions in 8D resulted in rapid disruption and loss of initial current inhibition, and a progressive increase of non-rectifying current. These results provide evidence that NT_2-4_, particularly I2/F2, in combination with M48 are primary determinants of activating vs. inhibitory current modulation by ChlT. M48 oxidation limits 8C inhibition and is required for activating responses, while T48 and 8D EL1 promote 8D signature current inhibition. ChlT exposure disrupts subsequent or preexisting channel block by DCPIB, consistent with a common site of interaction. Thus, factors that alter NT pore stability and mobility may regulate inhibition vs. activation of LRRC8C by redox stress.

## INTRODUCTION

Volume Regulated Anion Channels (VRACs), encoded by LRRC8 family genes (LRRC8A-E), have important physiological roles beyond regulation of cell volume (Thöne et al., 2025). Among these, LRRC8A (8A) (Choi et al., 2016) and LRRC8C (8C) (Choi et al., 2021) proteins physically associate with NADPH oxidases (Noxes) and support extracellular superoxide (O_2_^•-^) generation. 8A/C channels also provide an influx pathway for O_2_^•-^ (Harris et al., 2025; Panja et al., 2025), facilitating its delivery to intracellular targets associated with the channel LRR domains (Choi et al., 2023; Lamb et al., 2024; Harris et al., 2025). 8A/C channels are most closely associated with these pro-inflammatory functions. 8A or 8C knockdown in vascular smooth muscle cells (VSMCs) impaired O_2_^•-^ production and downstream TNFα signaling, while 8D knockdown had pro-inflammatory effects (Choi et al., 2023; Lamb et al., 2024; Harris et al., 2025). In accordance with these *in vitro* findings, VSMC-specific LRRC8A knockout (KO) mice displayed preserved vascular function when challenged by either AngII-induced hypertension (Choi et al., 2025) or ApoE KO-induced atherosclerosis (Panja et al., 2025).

Since VRACs regulate O_2_^•-^ production and influx, it is critical to understand how channel subtypes respond in an oxidized environment. Pro-inflammatory mediators including angiotensin II (Ren et al., 2008; Huo et al., 2021), H_2_O_2_ (Varela et al., 2004), and TNFα (Matsuda et al., 2010) have been reported to activate VRACs under isotonic conditions in a ROS-dependent manner, based on the impairment of activation by Nox inhibitors or ROS scavenging (Friard et al., 2021). However, LRRC8 channel subtypes exhibit variable current responses to the oxidant Chloramine T (ChlT). In oocytes and mammalian cellular expression systems, 8C and 8D channels activated by addition of C-terminal fluorescent tags or hypotonic saline display inhibitory current responses to ChlT and cysteine-modifying MTS reagents, while H_2_O_2_ and the membrane-permeant peroxide analog TBHP exert little effect (Gradogna et al., 2017b; Choi et al., 2021; Bertelli et al., 2022). 8A/E channels demonstrated current activation or augmentation in response to ChlT and TBHP (Gradogna et al., 2017b; Bertelli et al., 2022). This response was attributed to disulfide formation between cytoplasmic LRR domain cysteines unique to LRRC8E (Bertelli et al., 2022). A comprehensive mutational screening of redox-sensitive cysteines, methionines and histidines in 8C identified the start methionine (M1) as the principal target of inhibitory current modulation by ChlT (Bertelli et al., 2022). The insensitivity of 8C and 8D currents to H_2_O_2_ and TBHP underscores the question of how ChlT exerts its unique modulatory effects. We previously reported that currents mediated by homomeric 8D or heteromeric 8A/D channels were more potently inhibited by ChlT than 8C or 8A/C channels (Choi et al., 2021). Exchanging the 8C and 8D first extracellular loops (EL1) altered current responses to ChlT (Choi et al., 2021), suggesting that subtype-specific external targets largely account for this difference. However, subsequent clarification of TM1-EL1 domain boundaries showed that these substitutions (Yamada and Strange, 2018b) included the distal TM1_45-49_ region. 8C and 8D differ in TM1_45-49_ sequence and methionine position (M48 vs. M45), raising the possibility that the TM1_45-49_ pore region has a role in regulating the ChlT response.

Mutational studies have likewise demonstrated a critical role for the N-terminus (NT), particularly NT_1-15_, in LRRC8C/8A channel gating. The NTs form a critical part of the channel pore, and channel function is particularly sensitive to NT_2-4_ sequence and integrity (Zhou et al., 2018; Liu et al., 2023). Analyses of 8D and 8A channel structures have confirmed that the NT is extensively folded into the transmembrane portion of the channel pore, bringing NT_1-11_ into close proximity with the pore-lining TM1 helix (Nakamura et al., 2020; Liu et al., 2023). This is consistent with the accessibility of NT pore residues in 8A, 8C, and 8D channels to soluble cysteine-modifying agents (Zhou et al., 2018). The 8C NT pore structure has not been resolved, but likely more closely resembles the structure of 8A than 8D channels (Yamada et al., 2025). Compared to 8A, the 8D NT shows a less ordered structure and does not form protein-protein interactions with neighboring pore-forming structures, suggesting greater mobility within the pore (Liu et al., 2023). This is consistent with subtype-specific roles for the NT in regulating channel function (Zhou et al., 2018; Liu et al., 2023).

Recent work also demonstrates the presence of lipids closely associated with both external and internal surfaces of the LRRC8 pore, that play a critical role in channel gating (Kern et al., 2023) and binding of channel modulators (Yamada et al., 2025). Substitution of a negatively charged residue (T48D, M48D) at the apex of the TM1 helix resulted in 8A/C channel delipidation (Kern et al., 2023), and disrupted gating and reduced current inhibition by channel blockers in 8A/C and 8C channels (Kern et al., 2023; Yamada et al., 2025). These studies increasingly point to the role of the lipid-pore co-environment in regulating channel function.

We investigated chimeric 8C/8D channel constructs with the aim of identifying features of 8D responsible for conferring markedly stronger inhibitory ChlT sensitivity. We utilized C-terminal GFP-tagged channels substituted with the 8A intracellular loop (IL1) sequence (121 aa), which confers partial current activation under isotonic conditions (Gaitán-Peñas et al., 2016), while 8A IL1 traffics these homomeric channels to the plasma membrane (Yamada and Strange, 2018a; Choi et al., 2021). We show that 8D NT_2-4_ substitutions confer current-activating responses in 8C channels in response to ChlT. This response requires the presence of M48. T48 substitution in 8C enhances ChlT-dependent current inhibition, while M48 substitution in 8D disrupts inhibition and activates an atypical non-rectifying current, indicating that M48 oxidation limits current inhibition and is necessary for current activation. This form of oxidative current modulation is thus sensitive to NT and TM1 protein structure, and to lipid interactions with pore amino acids. Physiological regulation of NT mobility may therefore influence how LRRC8C oxidation affects channel activity and modulates pro-inflammatory signaling.

## MATERIALS AND METHODS

### Reagents and Cell Culture

All chemicals and enzymes were obtained from Sigma-Aldrich (St. Louis, MO) unless otherwise noted. LRRC8A knockout (LRRC8A^(*-/-*)^) HEK293T cells were a gift from Dr. Rajan Sah (Washington University, St. Louis MO). Cells were maintained at 37° C in 5% CO_2_ in Dulbecco’s modified Eagle’s medium (DMEM) supplemented with 10% fetal bovine serum (FBS, Hyclone) and 0.25% Pen/Strep (25U/ml, Life Technologies).

### Plasmid Modification and Expression

LRRC8C and LRRC8D constructs substituted with the LRRC8A intracellular loop (IL) (Choi et al., 2021) were provided by Dr. Kevin Strange (Novo Biosciences and Vanderbilt University). These constructs, bearing C-terminal GFP tags, are referred to here as wild-type 8C and wild-type 8D. Plasmid modifications were made using the QuikChange Lightning site-directed mutagenesis kit (Agilent). Plasmids were transfected using Lipofectamine 2000 (DMEM + 1.5-2.5 µg/µl cDNA, 2.5% lipofectamine 2000, +10% FBS). Transfection media was replaced with fresh DMEM after 6-18 hrs.

### Electrophysiology and Cell Imagining

LRRC8A^(*-/-*)^ HEK293T cells were seeded into 6- or 12-well plates and allowed to adhere for either 4-6 hrs or overnight prior to transfection. Whole-cell patch clamp recordings were made at room temperature (21-22 °C) from freshly trypsinized and dissociated cells on the 1^st^ (18-24 hrs) or 2^nd^ day (42-48 hrs) post transfection. Protein expression and distribution was documented in recorded cells by visualizing fluorescence on an Olympus IX71 microscope (60X objective) equipped with a Hamamatsu C10600 digital camera and Till Photonics Oligochrome and image acquisition/analysis hardware and software (Hunt Optics & Imaging, Inc, Pittsburgh, PA). Isotonic external recording saline (330 mosm/L, pH 7.35) contained (in mM): 130 NaCl, 1.8 MgCl_2_, 1.8 CaCl_2_, 10 HEPES, 70-75 Mannitol, 1-2 NaOH. Pipette saline (315 mosm/L, pH 7.2) contained (in mM): 120 CsCl, 4 TEACl, 2 MgCl_2_, 5 Na_2_ATP, 10 HEPES, ∼35 Mannitol, 1-2 CsOH, 1.186 CaCl_2_, 5 EGTA (estimated free [Ca^2+^] of 61 nM using WEBMAXC). Final solution osmolality was measured with a Precision Systems mOsmette osmometer (Natick, MA) and adjusted by addition of 1M mannitol. External salines containing Chloramine T (ChlT, 1.0 mM), MTSET (1.0 mM), hydrogen peroxide (H_2_O_2_, 0.5 mM), TBHP (1.0 mM), sodium hyperchlorite (NaOCl, 1 mM), and dithiothreitol (DTT, 5 mM) were freshly prepared daily. 0.5-1.0 M stock solutions (ChlT, MTSET) were freshly remade weekly. DCPIB (30 μM; Cayman Chemical, Ann Arbor MI) was freshly added from 30 mM (DMSO) stock aliquots stored at −20 °C. The cell recording chamber was perfused at 1.2-1.5 ml/min and maintained at a volume of 0.4-0.5 ml via an Automate ValveBankII system (Berkeley, CA). Saline changes and drug applications were controlled via a multiport manifold that delivered new solution to the chamber within 7-8 sec. DCPIB and ChlT effects on current were typically observable within 20-30 sec after toggling to drug-containing saline.

Amplified whole-cell currents were low-pass filtered at 5 kHz and sampled at 5-10 kHz using a Molecular Devices Axopatch 200B amplifier driven with a pClamp 10 interface (Molecular Devices, Sunnyvale, CA). Pipette resistance after fire-polishing and filling with pipette solution was 2.0-3.0 MΩ. Pipette capacitance was nulled following gigaseal formation. Cell capacitance (22.7 ± 0.7 pF) and series resistance (Rs; 5.4 ± 0.2 MΩ) were measured after establishing whole cell configuration using the Clampex 10 membrane test utility at a holding potential (V_H_) of −40 mV. 75%-90% R_S_ correction (>90% prediction) was applied. R_S_ was monitored at intervals throughout recordings and correction was readjusted as necessary. VRAC currents were elicited with 250 ms voltage ramps (−100 to +120 mV) beginning ∼2 min after establishing whole cell configuration. Baseline current was recorded for 2’-5’ until amplitude reached stability.

Standard current-voltage (I-V) relationships were recorded in a subset of cells using 100 ms test pulses (−100 to +120 mV, 20 mV increments) applied from a prepulse level of 0 mV. Current amplitude was measured 2 ms after test pulse onset. Time-dependent current changes and drug effects were assessed with ramp-evoked current responses recorded at 20s intervals and quantified at −100 and +120 mV. Current density is expressed as pA/pF, and I-V relationships are corrected for a +5 mV liquid junction potential measured between the pipette and isotonic bath saline. Zero-current level is indicated by a dashed line in displayed traces.

### Statistical analysis

Statistical calculations and comparisons were made with Excel (2013) and Graphpad Prism 10 analysis software (GraphPad Software, San Diego, CA). Exported current traces (refiltered at 1 kHz) and plots were generated and displayed using GraphPad Prism. Values are reported as Mean ± S.E.M. unless otherwise indicated. N represents the number of recorded cells for the indicated data group. Ratio T-tests were applied to log-transformed current amplitudes (+120 mV and −100 mV) to compare paired control and drug-treated measurements. Wilcoxon paired tests or Mann-Whitney nonparametric tests were used for other two-way comparisons. Non-parametric ANOVA with Dunn’s test was used for multiple comparisons. *P* values less than 0.05 were considered statistically significant.

## RESULTS

### Isotonic currents of LRRC8C and LRRC88D channels are differentially inhibited by ChlT

Chimeric LRRC8C and LRRC8D proteins substituted with the 8A first intracellular loop sequence form functional homomeric channels in the plasma membrane of LRRC8A^(-/-)^ HEK293 cells that lack functional native LRRC8 channels (Yamada and Strange, 2018a; Choi et al., 2021). C-terminal eGFP tags were added to confer partial channel activation under isotonic conditions (Gaitán-Peñas et al., 2016; Gradogna et al., 2017a; Yamada and Strange, 2018a; Choi et al., 2021). We refer to these channels here as wild-type (WT) 8C and 8D. Previously, we documented inhibitory effects of ChlT on 8C and 8D channel currents activated by hypotonic conditions, and modulation of inhibition by the extracellular loop (EL) domains (Choi et al., 2021). Recent structural analyses (Nakamura et al., 2020; Liu et al., 2023; Rutz et al., 2023; Quinodoz et al., 2025) have clarified that residues 45-49 previously mapped to EL1 (Yamada and Strange, 2018a) make up the distal transmembrane (TM1) helix (Fig. 1), raising the possibility that TM1_45-49_ has a role in ChlT-dependent current modulation. We addressed the action of ChlT and other redox agents on 8C and 8D channel currents under isotonic conditions, focusing on the pore-lining N-terminal (NT) and distal TM1.

**Figure 1.**
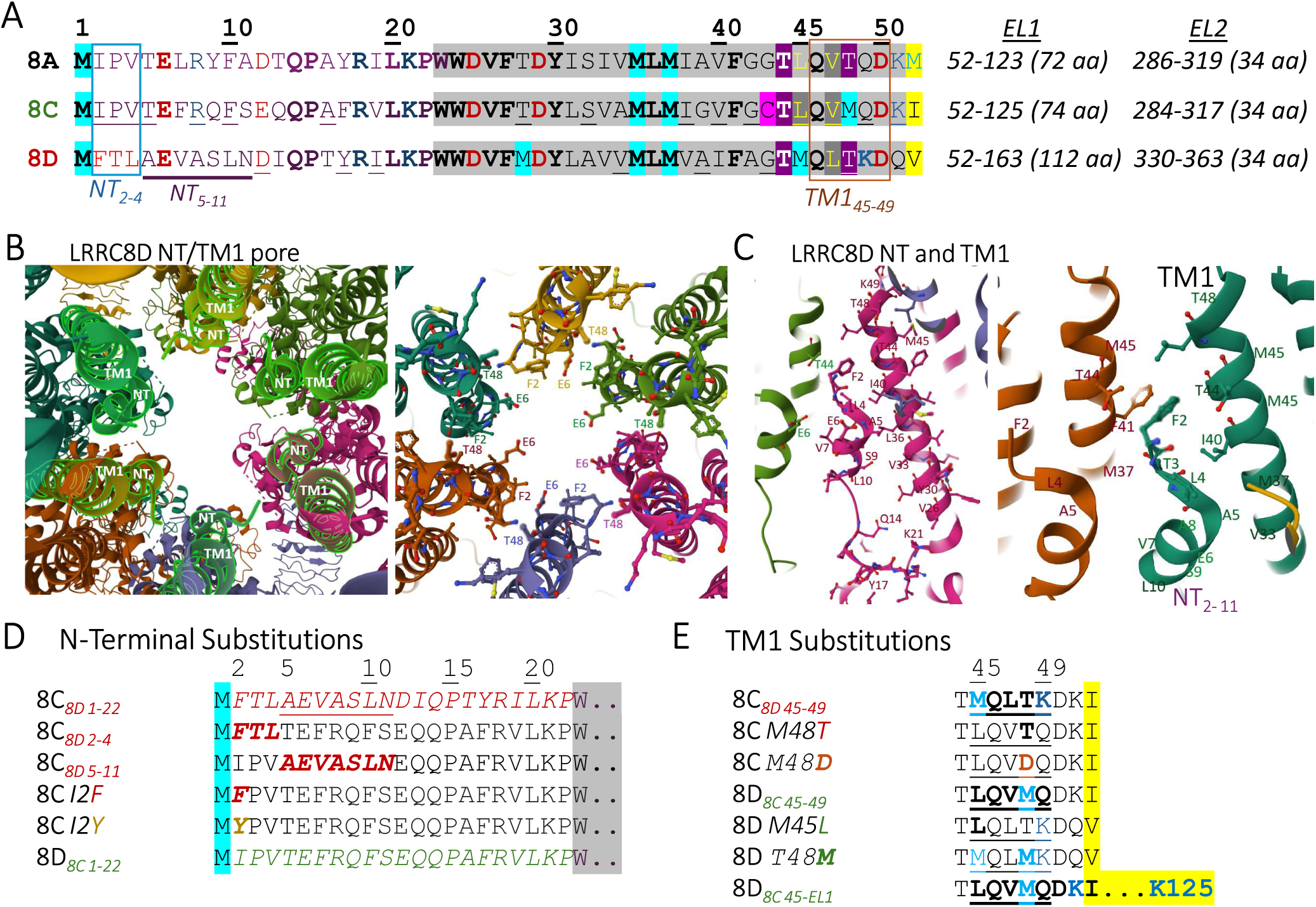
LRRC8C and 8D N-terminal and TM1 sequences and substitutions. ***A***, Comparison of N-terminal (aa 1-22) and TM1 (aa 23-49 gray shading) protein sequences of LRRC8A, 8C, and 8D channels. Conserved residues are shown in bold; common residues between 8C and 8A and the 8D NT_2-11_ region are underlined. NT_2-4_ and TM1_45-49_ regions are outlined. The beginning of the first extracellular loop domain (EL1) is shaded in yellow. Potential targets for oxidation (M, teal; C, pink) are indicated. ***B***, Two external views of the LRRC8D channel viewed at 90° to the membrane. ***Left***, the six TM1 and NT regions surrounding the channel pore (green outline) are highlighted. ***Right***, TM1- and NT-lined pore showing F2, E6, and T48 sidechains. M1 is not represented. ***C, Left.*** Vertically oriented view of an individual 8D NT-TM1 structure, showing NT_b_ helix (L4-L10) opposed to the TM1 helix. ***C, Right***, View of 8D NT_2-11_ and distal TM1 (33-48) regions in adjacent subunits. F2 extends between I40 and T44. Structures cited from PDB 6M04. The NT has not been resolved in reported 8C structures (not shown). ***D***, 8C and 8D NT sequence variants. Substituted residues (8D 1-22, 2-4, 5-11, I2F, and I2Y) are italicized. Gray shading indicates the beginning of TM1. ***E***, 8C and 8D TM1_45-49_ sequence variants, with substituted residues shown in bold. The 8C_45-EL1_ substitution (L45-K125; bottom) replaced the entire 8D TM1_45-49_ + 8D EL1 sequence.

The LRRC8A NT (aa 1-22) and TM1 (aa 23-49) sequence shares 73% identity with 8C, but only 57% with 8D (Fig. 1A). The three sequences are highly similar from Q14-F41, but differ notably within the NT_2-11_ and TM1_45-49_ regions, where 8A and 8C share 11/15 identical amino acids, but 8D shares only 3/15 identical residues with 8A and 2/15 with 8C (Fig. 1A). 8A and 8D channel structures with resolved NTs show the NT to be folded extensively into the channel pore, extending most of the transmembrane distance and opposed to the TM1 helix and neighboring NT (Nakamura et al., 2020; Liu et al., 2023) (Fig. 1B, C). Mutational studies have shown that NT sequence regulates 8A/C channel current and current modulation by oxidants (Zhou et al., 2018). To examine subunit-dependent roles for the NT in 8C and 8D current modulation, we tested a series of related 8D-to-8C NT sequence substitutions (Fig. 1D). We then tested subunit-specific substitutions within the TM1_45-49_ region of 8C (LQVMQ) and 8D (MQLTK) to assess the potential for methionine-dependent current modulation.

Cells expressing WT 8C exhibited 2-fold larger basal isotonic current amplitudes overall, and significantly stronger current rectification than in 8D-expressing cells (P < 0.0005 vs 8D; Fig. 2Af, 2Bf, and Table 1). 1 mM ChlT was applied for 3-5’ while recording current responses to voltage ramps (−100 to +120 mV) at 20s intervals. 8C currents were moderately but significantly inhibited by ChlT (30 ± 6 % inhibition vs. Iso control, P < 0.0005; n = 10) (Fig 2Ab-f). The 8D current response was distinguished by more rapid onset of inhibition (20-30s), and substantially more potent overall inhibition (85 ± 3 % inhibition vs. Iso control, n = 14; P < 0.0005 vs 8C WT) (Fig. 2Bb-f). Among individual recordings, the degree of ChlT-mediated inhibition (I_ChlT_/I_Con_) was uncorrelated with basal control current amplitude (Supplementary Fig. S1). For both 8C and 8D, inhibition of outward current (inward anion flux) was more pronounced than for inward current (Fig. 2A, B, panel f). As reported previously (Choi et al., 2021), addition of 30 μM DCPIB blocked 8C and 8C currents by 90-95% Fig. 2A,B, panel a).

**Figure 2.**
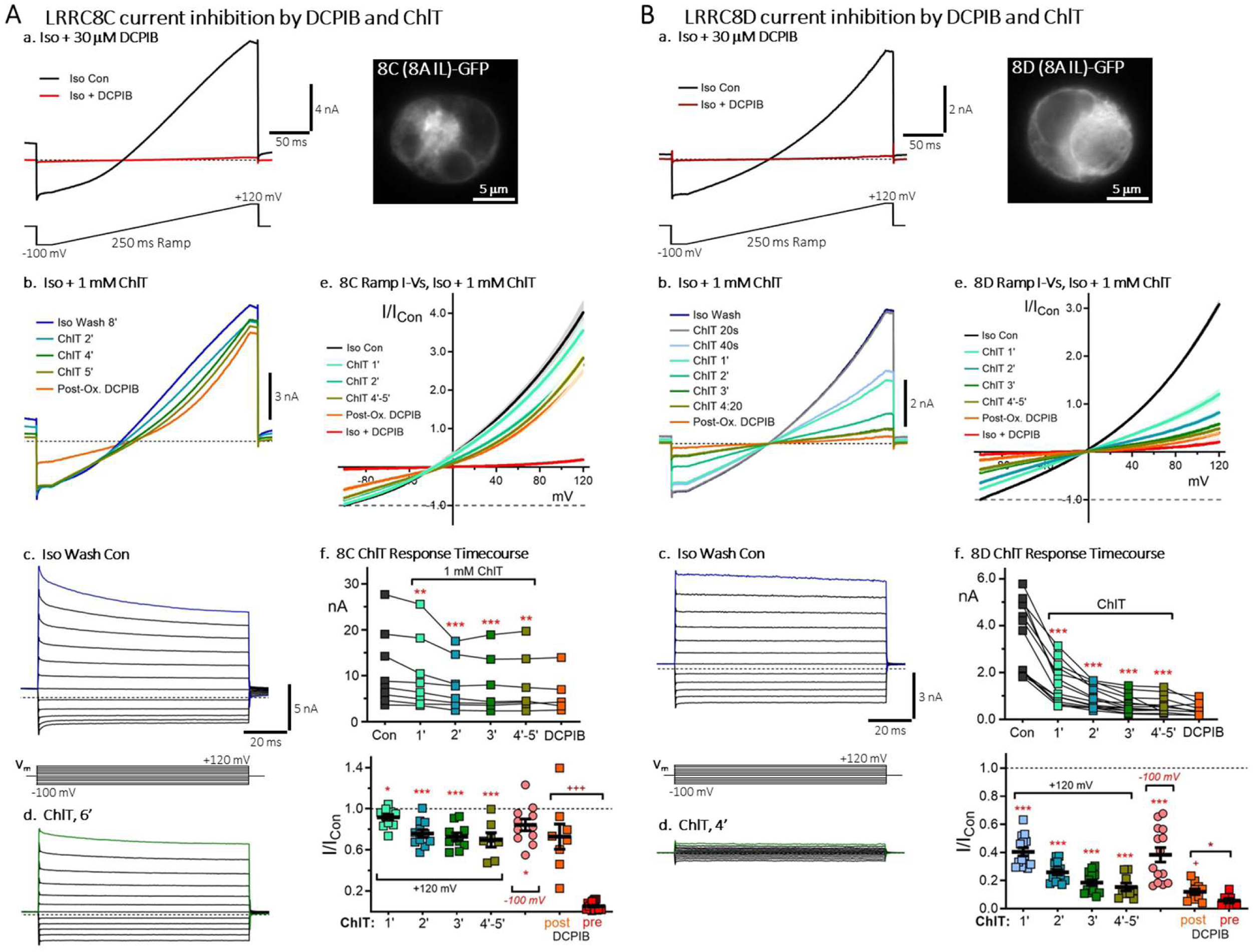
Wild-type LRRC8C and LRRC8D channels display distinctive current inhibition by Chloramine-T (ChlT). Representative currents recorded in LRRC8A*^(-/-)^*HEK cells expressing “wild-type” GFP-tagged LRRC8C (***A***) or LRRC8D (***B***) chimeric proteins substituted with the LRRC8A intracellular loop sequence (*8A IL*). ***Panel a,*** Rectifying currents elicited by 250 ms voltage ramps (−100 to +120 mV; ramp depicted at bottom) in isotonic control saline (Iso Con). Control currents are blocked by 90-95% by 30 μM DCPIB (red traces; dotted lines indicate zero-current level). Cell images (***right***) illustrate protein expression in the recorded cells (scale: 5 μm). ***Panel b,*** Currents recorded in the same cells shown in (**a**) following DCPIB washout (Iso Wash). 8C currents are moderately inhibited by 1 mM ChlT (***Ab***), while 8D currents are more rapidly (20 s - 3’) and potently inhibited (***Bb***). ***Panels c-d,*** Currents elicited by 100 ms voltage steps (−100 to +120 mV) in Iso Con (***c***) and following ChlT exposure (***d***); Dotted lines indicate zero-current level. ***Panel e***, Mean ramp current-voltage (I-V) relationships (± S.E.M.) for Iso control and ChlT-treated conditions, normalized to control current amplitudes at −100 mV (dotted line). Plots illustrate differential inhibition by 1 mM ChlT between 8C (n = 10) and 8D (n = 14), as well as a loss of DCPIB sensitivity following ChlT exposure (Post-Ox. DCPIB; orange traces). ***Panel f***, Time course of current amplitude changes (nA, +120 mV; ***top***) and normalized current (I/I_Con_, ***bottom***) during 4’-5’ ChlT treatment in individual recordings. 8C-expressing cells exhibited larger basal current amplitudes. ***Bottom***: Values at −100 mV are included (4’-5’ ChlT; pink symbols). Post-ChlT DCPIB values (orange symbols) are compared to pre-Ox. DCPIB current (red symbols). Bars show mean values (± S.E.M). Asterisks indicate statistical significance compared to Iso control values, or between indicated pairs (*** P < 0.0005; *** P < 0.005; * P < 0.05).

**Table 1.**
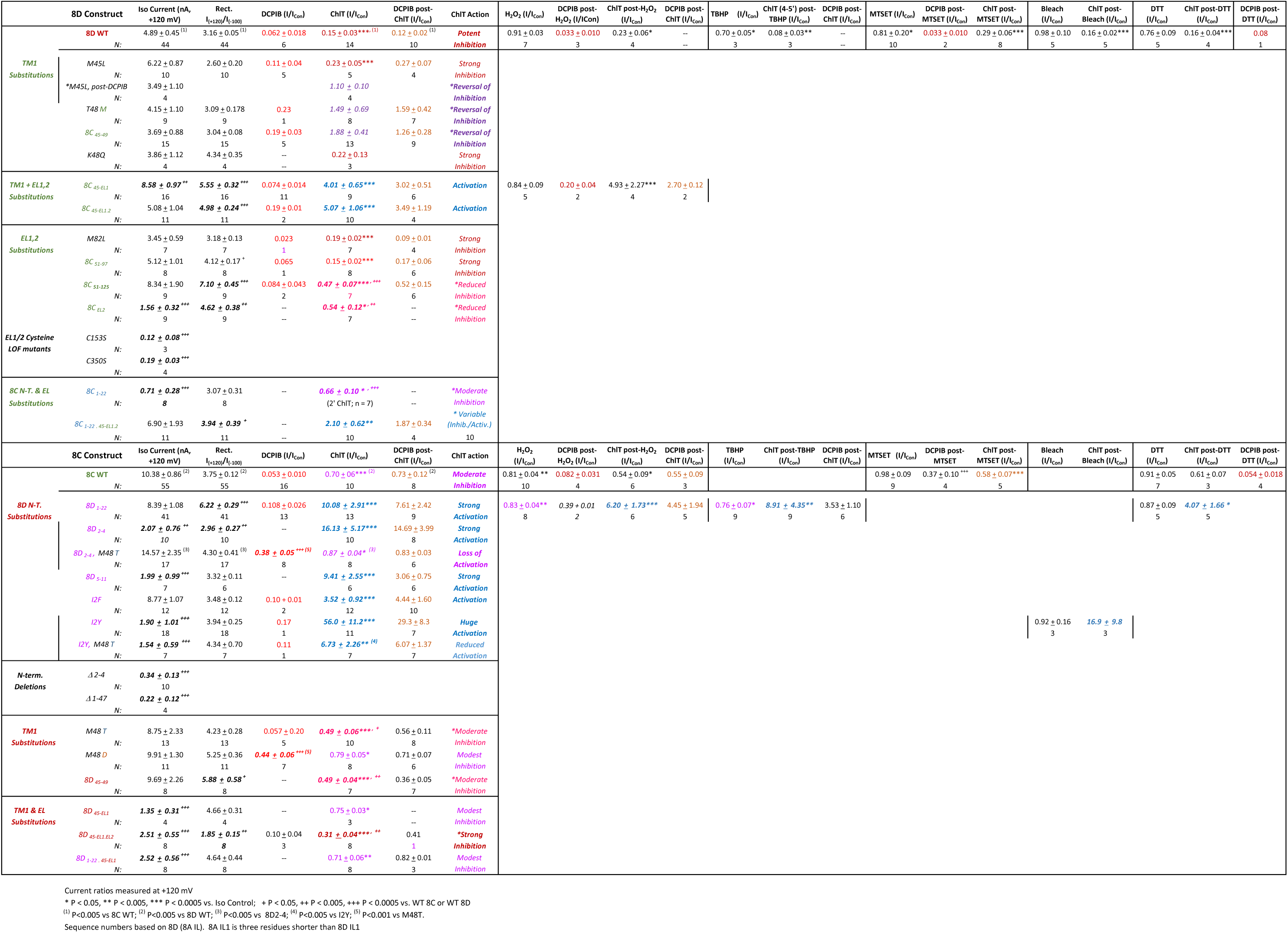

However, following ChlT exposure, current block by DCPIB was significantly reduced or absent (“post-ox.” DCPIB; Fig. 2A,B, panels e-f and Table 1). This was especially evident for 8C, which had large post-ChlT currents that showed no overall inhibition by DCPIB (n = 7; Fig 2Af).

### ChlT uniquely modulates 8C and 8C channel currents and channel block

Both cytoplasmic and extracellular methionine and cysteine targets have been proposed to regulate the inhibitory action of ChlT on 8C and 8D currents (Gradogna et al., 2017b; Choi et al., 2021; Bertelli et al., 2022). Other studies primarily examined 8C/8A and 8D/8A heteromeric channels in oocytes and human cellular expression systems, with ChlT the principal oxidant examined. ChlT hydrolyses in solution to form hypochlorous acid (HOCl), making it an effective chlorinating as well as oxidizing agent. To compare the action of ChlT to other redox active agents on homomeric 8C and 8D channels, we examined current responses to H_2_O_2_ (0.5 mM) and the lipid-permeable tert-butyl hydroperoxide (TBHP; 1 mM), a cell-impermeable cysteine modifier MTSET (1 mM), sodium hypochlorite (NaOCl; 1 mM), and the reducing agent DTT (5 mM) (Fig. 3). Surprisingly, 8D (Fig. 3A) and 8C (Fig. 3B) currents are comparatively insensitive to most of these agents. 8D currents were modestly inhibited by MTSET (19 ± 20 % inhibition, n = 10; Fig. 3Aa) and TBHP (30 ± 5 % inhibition, n = 3: Fig. 3Ac) (P < 0.05 vs. Iso Con). 8C currents were likewise moderately inhibited by H_2_O_2_ (19 ± 4 % inhibition, n =10; Fig. 3Bb; P < 0.005 vs. Iso Con). Currents were not significantly modified by DTT. The efficacy of current block by DCPIB was not significantly impaired by exposure to these other agents (Fig. 3C, D), unlike the effect of ChlT treatment, with the exception MTSET which reduced post-ox. DCPIB block of 8C current (P < 0.0005 vs Iso DCPIB control; Fig. 3D). Additionally, none of the other agents prevented or masked the subsequent inhibitory action of ChlT on 8C or 8D currents shown in Fig. 2 (Fig. 3C, D). The unique actions of ChlT on both current and channel block indicate that ChlT acts on exclusive target(s) or possesses a unique potency for oxidative current modulation.

**Figure 3.**
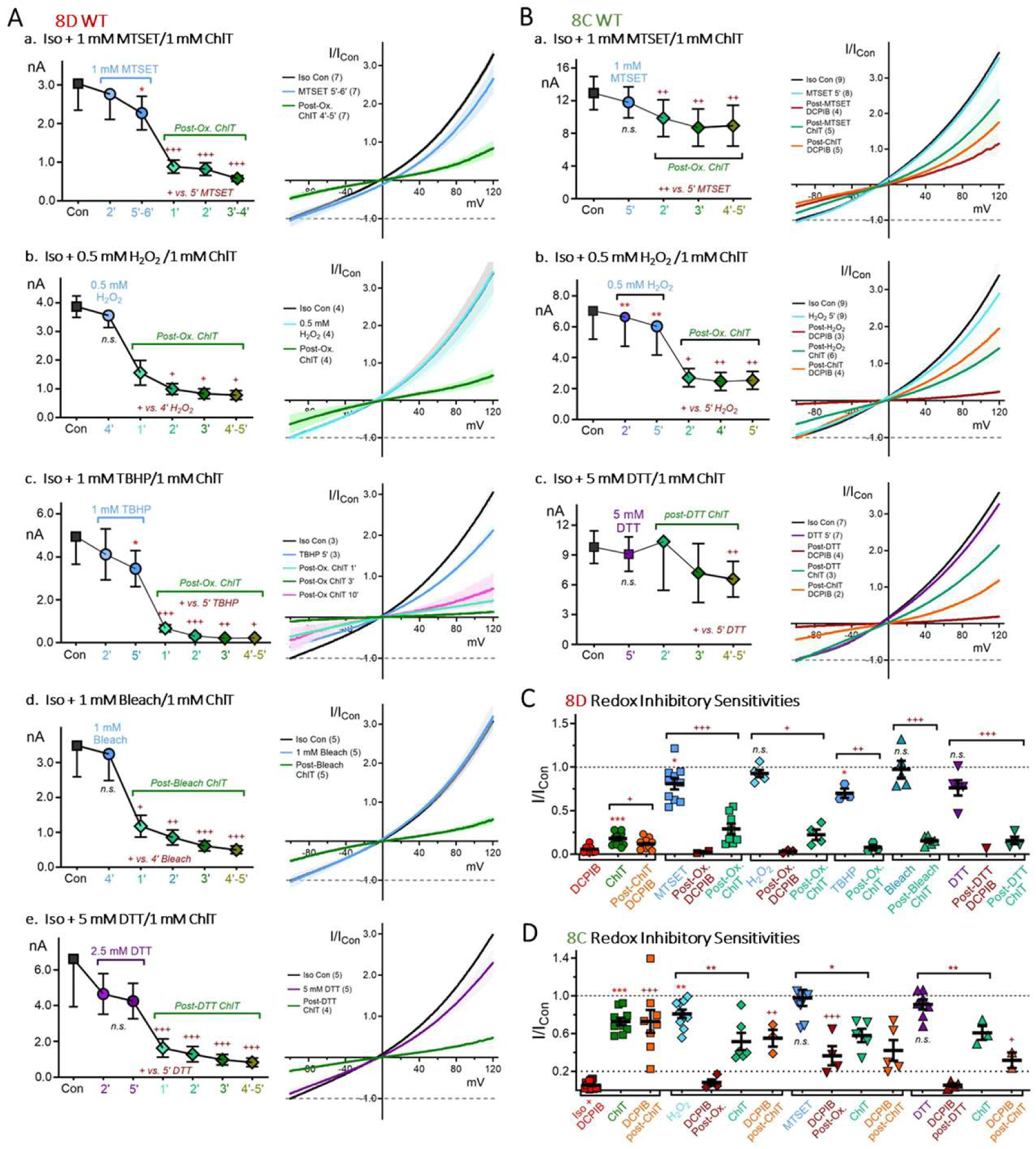
LRRC8C and 8D channel currents are less sensitive to other redox agents compared to ChlT. ***A,*** 8D isotonic currents. ***a-e, left***: Plots show time course of mean (± S.E.M.) current amplitude (+120 mV) changes during 4’-5’ application of 1 mM MTSET (***a***), 0.5 mM H_2_O_2_ (***b***); 1 mM TBHP (***c***); 1 mM Na-Hypochlorite (bleach, ***d***); 5 mM DTT (***e***), followed by application of 1 mM ChlT (green symbols). 8D current is weakly (10-15%) inhibited by MTSET (***a***) and TBHP (***c***). Current decrease produced by DTT was not significant. In all cases, subsequent exposure to ChlT produces the expected degree of current inhibition. Asterisks indicate significance change vs. Iso control (*, P < 0.05). For ChlT, symbols indicate significance vs. 5’ initial drug treatment (^+^, P < 0.05, ^++^ < 0.005, ^+++^ < 0.0005). ***a-e, right***: Mean I-V relationships for control and drug-treated conditions, normalized to control current at −100 mV (dotted line). ***B***, 8C isotonic currents. ***a-c, left***: Time course of mean current amplitudes (+120 mV), as in ***A***. 8C current is weakly inhibited by 0.5 mM H_2_O_2_ (***b***), but insensitive to 1 mM MTSET (***a***) and 5 mM DTT (***c***). In each condition, subsequent ChlT exposure produces significant inhibition (significance indicated as in ***A***). ***a-c***, ***right***: Mean I-V relationships for control and drug-treated conditions. 8C currents remain sensitive to DCPIB block (80-90% inhibition) following H_2_O_2_ and DTT treatment (magenta traces, ***b-c***), but show impaired DCPIB sensitivity following MTSET (***a***, 50%-80% inhibition). DCPIB block is strongly diminished after ChlT exposure (***a-c***, orange traces). ***C-D***, Summary of 8D (***C***) and 8C (***D***) current response to redox agents and DCPIB (+120 mV; *, P < 0.05, ** < 0.005, *** < 0.0005 vs. Iso control). Crosses (+, ++, +++) indicate comparison of ChlT current to pre-ChlT drug treatment (horizontal brackets). In ***D***, post-drug DCPIB current is compared to pre-drug block level (Iso + DCPIB control).

### 8D NT substitution confers activating responses to 8C

The NT-resolved LRRC8A and 8D channel structures show the NT to be integral to the channel pore and positioned to interact with the distal TM1 helix protein and/or lipid environment. 8D (M45) and 8C (M48) have potentially reactive methionines in distinct TM1 positions, in addition to the NT M1. The 8D NT_2-4_ sequence (FTL) is unique among LRRC8 proteins, raising the possibility of an 8D-specific regulatory role. As an initial step to assessing the role of the NT in regulating channel oxidant sensitivity, the entire 8D NT_1-22_ sequence was substituted into 8C. Currents in 8C*_8D 1-22_*-expressing cells had robust amplitudes similar to WT 8C, but exhibited stronger rectification (P < 0.001 vs WT 8C; Table 1 and Supplementary Fig. S2). Remarkably, 8C*_8D 1-22_* current amplitudes increased in response to ChlT (Fig. 4A), which often led to amplifier-saturating current amplitudes in initial recordings. We consequently biased selection toward cells with weaker GFP expression levels for most subsequent recordings, resulting in smaller basal current amplitudes (2.21 ± 0.64 nA; n = 13). This approach clearly revealed that current activation (I/I_Con_ at +120 mV) was inversely related to initial current level (see below), with ChlT eliciting up to 30-fold increases in current amplitude, and 10-fold overall activation (Fig. 4Ae-g; Table 1). The onset of current activation was slightly delayed compared to WT inhibitory responses, but activation was clearly evident by 1-1.5’ after ChlT exposure, and peaked after 4-6’ (Fig 4A a,b,e,f). 8C*_8D 1-22_* currents were strongly blocked by DCPIB application in normal saline (89 ± 3 % block; n =13), but as in WT post-ChlT current block by DCPIB was nearly eliminated (Fig. 4A, e-g).

**Figure 4.**
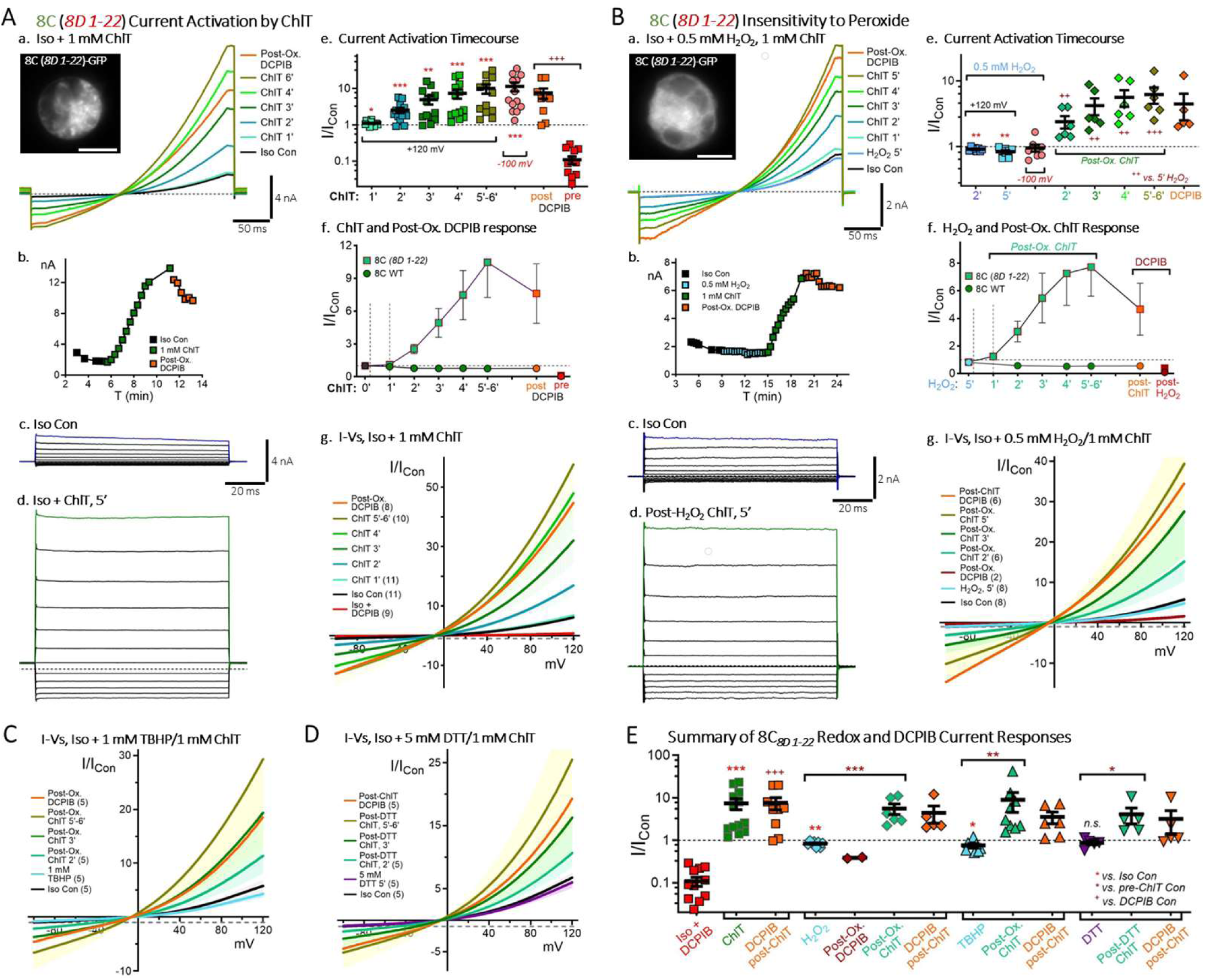
LRRC8D N-terminus substitution in LRRC8C confers current-activating responses to ChlT. Images (***A, B top left***) show cellular distribution of 8C*_8D 1-22_* protein in the recorded cells (scale: 5 μm). ***Aa,*** Ramp-generated (−100 to +120 mV) 8C*_8D 1-22_* currents in isotonic saline are dramatically activated from basal control levels during 5’-6’ exposure to 1 mM ChlT. ***b***, Time course of current amplitudes (+120 mV) recorded in the same cell. Current activation is evident within 1’ after ChlT application. ***c-d***, Currents activated by 100 ms voltage steps (−100 to +120 mV) under control conditions (***c***) and following 5’ ChlT exposure (***d***) in the same cell as ***a-b.*** Dotted lines indicate zero-current level. ***e,*** Time course of normalized current amplitudes (I/I_Con_, +120 mV) in individual recordings during ChlT (green, blue and brown symbols) and post-ox. DCPIB application (orange symbols). Values at −100 mV at 4’-5’ are included (pink symbols). Bars show mean values ± S.E.M. Asterisks indicate statistical significance compared to Iso control values, or between indicated group pairs (*** P < 0.0005; ** P < 0.005; * P < 0.05). ***f***, Normalized current amplitude time course (I/I_Con_, +120; mean ± S.E.M.) illustrating overall activation by ChlT, and almost complete loss of DCPIB sensitivity following ChlT exposure (post-ox. DCPIB, orange symbols), compared to pre-ChlT DCPIB block (red symbols). The WT 8C response is included for comparison (circles). ***g***, Mean I-V relationships (± S.E.M.) for 8C*_8D 1-22_*, normalized to Iso control current at −100 mV (dotted line). ***Ba, b,*** 8C*_8D 1-22_* currents are marginally inhibited by 5’ application of 0.5 mM hydrogen peroxide (H_2_O_2_, light blue trace/symbols). Subsequent application of 1 mM ChlT results in pronounced current activation ***b***, Time course of current amplitudes (+120 mV) in the same same cell. ***c-d***, Currents activated by 100 ms voltage steps (−100 to +120 mV) under control conditions (***c***), and following 5’ ChlT exposure post-H_2_O_2_ (***d***) in the same cell shown in ***a-b***. Dotted lines indicate zero-current level. ***e,*** Time course of normalized current amplitudes (+120 mV) in individual recordings during H_2_O_2_ (blue symbols), DCPIB (red and orange symbols), and post-H_2_O_2_ ChlT. Significance is indicated as in ***Ae***. For ChlT, symbols indicate significance vs. 5’ H_2_O_2_ treatment (+++, P < 0.0005; **, ++ P < 0.005). ***f***, Normalized current amplitude time course (I/I_Con_, +120 mV; mean ± S.E.M.), illustrating ∼10% overall inhibition by H_2_O_2_, in contrast to strong subsequent current activation by ChlT. The WT 8C response is included for comparison (circles). DCPIB produces substantial current inhibition following H_2_O_2_ (dark red symbols; n = 2), but DCPIB block is largely lost following subsequent ChlT exposure (orange symbols; n = 5). ***g***, Mean H_2_O_2_ and post-ox. ChlT I-V relationships (± S.E.M.) for 8C*_8D 1-22_*, normalized to control current at −100 mV (dotted line). ***C, D***, Mean TBHP and DTT I-V relationships (± S.E.M.). 8C*_8D 1-22_* currents are weakly (∼10%) inhibited by 1 mM TBHP (5’, light blue trace; ***C***) and insensitive to 5 mM DTT (5’ purple trace; ***D***). Subsequent ChlT exposure elicits strong current activation in both conditions. ***E***, Summary of 8C*_8D 1-22_* normalized current responses to redox agents and DCPIB. ChlT uniquely mediates strong current activation, in contrast to H_2_O_2_, TBHP, and DTT, which generate minimal or insignificant inhibitory responses. Asterisks indicate significance vs. Iso Con, or between indicated data pairs (horizontal arrows; *** P < 0.0005; **, P < 0.005; * P < 0.05). Crosses indicate significance vs. pre-ox and DCPIB.

The 8C*_8D 1-22_* current response to ChlT suggested that this background might confer altered sensitivity to other oxidants tested above. Contrary to this expectation, 8C*_8D 1-22_* currents were only weakly inhibited by H_2_O_2_ (P < 0.005 vs. control, n = 8; Fig. 4Ba,b,e,f) and TBHP (P < 0.05 vs. control, n = 9; Fig. 4E), while in both cases subsequent “post-ox.” ChlT application resulted in strong current activation (Fig. 4Be-g, Fig. 4E). 8C*_8D 1-22_* currents were also unaffected by DTT, and subsequently activated by ChlT exposure (Fig. 4E). ChlT consistently reduced or abolished subsequent current block by DCPIB (Fig. 4E). The “out-of-context” 8C*_8D 1-22_* substitution thus reversed the modulatory action of ChlT from inhibitory to current-activating, without significantly impacting current sensitivity to other redox agents.

### 8D NT_2-4_ substitutions are sufficient to confer activating responses to 8C

We next investigated whether substitutions from 8D NT_2-11_ sequence are able to confer current-activating responses to 8C. 8D_2-4_ substitution resulted in smaller initial currents (2.07 ± 0.76 nA, n = 10) with reduced rectification (P < 0.005 vs 8C WT; Fig. 5E, Supplementary Fig. S2 and Table 1). 8D_2-4_ currents were strongly increased by 10-15 fold in response to 1 mM ChlT (Fig. 5A, G). 8D_5-11_ substitution likewise resulted in reduced basal current amplitude (1.99 ± 0.99 nA, n = 6; P < 0.001 vs. WT 8C) that were activated by over 9-fold by ChlT (Fig. 5E-G). Deletion of 8C NT_2-4_ (Δ2-4) resulted in a complete loss of functional current, as previously reported (Zhou et al, 2018), as did a larger Δ1-47 deletion (Fig. 5E and Table 1).

**Figure 5.**
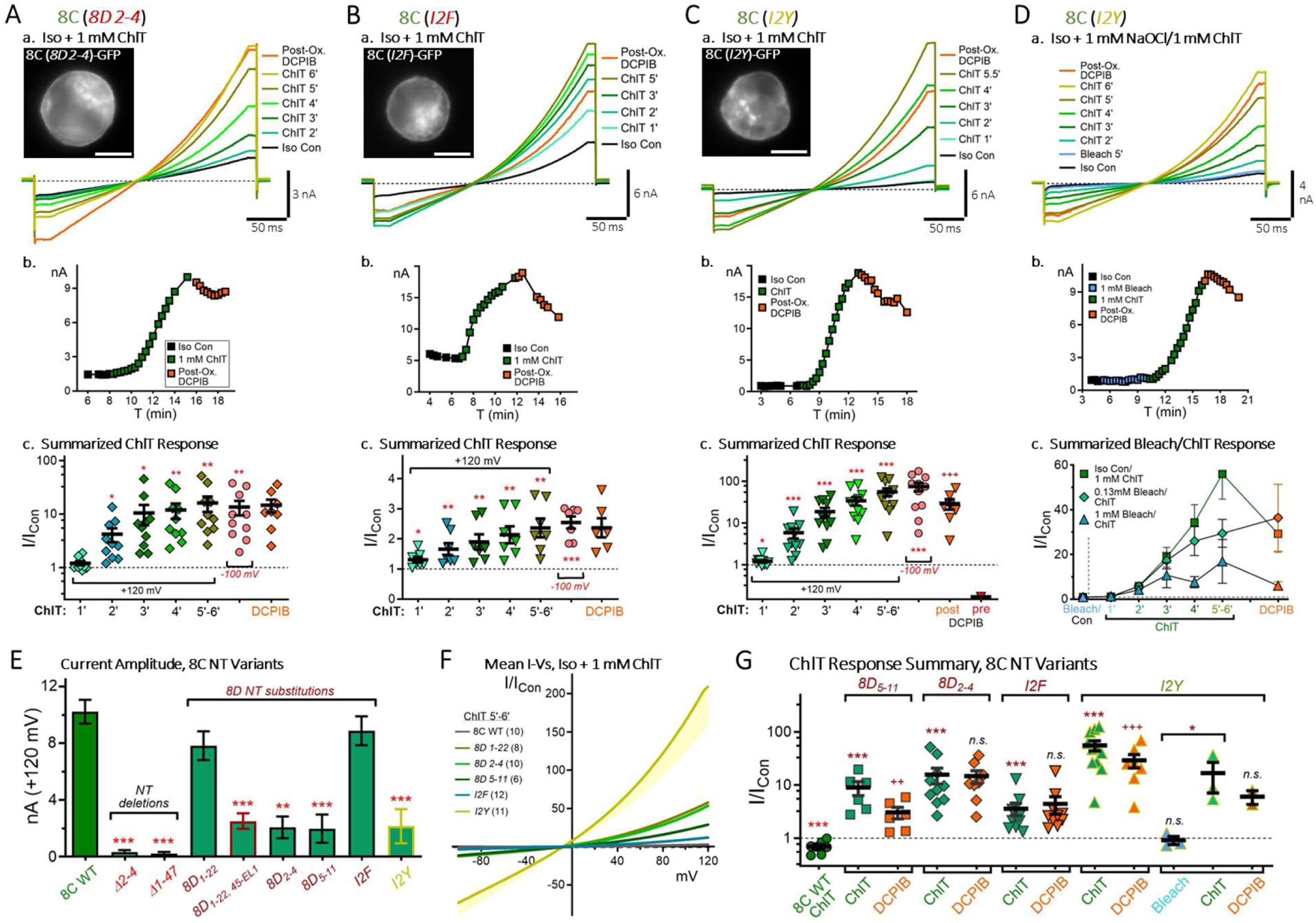
LRRC8D *NT_2-4_* substitutions in LRRC8C are sufficient to confer current-activating ChlT responses. ***A-D***, ***Panel a***, Ramp generated isotonic currents (−100 to +120 mV) in cells expressing LRRC8C with the indicated N-terminal substitution (***A***, *8D_2-4_; **B***, *I2F; **C, D***, *I2Y*). Images (***left***) illustrate protein distribution in the recorded cell (scale: 5 μm). Currents are activated by application of 1 mM ChlT (5’-6’). Dotted lines show zero-current level. ***Panel b***, Time course of current amplitude changes (+120 mV) for the recording shown in ***a***. ***Panel c, A-C***, Normalized current amplitude changes (I/I_Con_, +120 mV) over 5’-6’ exposure to ChlT (horizontal bracket) and post-ox. DCPIB (orange symbols) in individual recordings. Values at - 100 mV (4’-5’ ChlT, pink symbols) are included for comparison. Bars show mean values (± S.E.M.). Asterisks indicate significance compared to Iso control current (*** P < 0.0005; ** P < 0.005; * P < 0.05); crosses (***Cc***) indicate significance vs. 5’6’ ChlT (++ P < 0.005). ***D, a-c***: *I2Y* currents are unaltered by 1 mM bleach (light blue trace/symbols), but dramatically activated by subsequent application of 1 mM ChlT (green symbols). ***Dc***, Normalized mean current amplitude changes (I/I_Con_, +120 mV) to 5’ bleach exposure (light blue symbols) and subsequent exposure to ChlT (horizontal bracket) and post-ox. DCPIB (orange symbols; * P < 0.05 vs. Iso Control). ***E***, Isotonic basal current amplitudes for 8C N-terminal variants, including N-term deletions and 8D substitutions (*** P < 0.0005 vs. 8C WT). ***F***, Mean I-V relationships (± S.E.M.) for 8C (*8D_1-22_*)-, (*8D_2-4_*)-, (*8D_5-11_*)-, *I2F*-, and *I2Y*-expressing cells exposed to ChlT (1 mM, 5’-6’), normalized to Iso control current at −100 mV (dotted line). 8C WT trace is included for comparison. ***G***, Summary of normalized current responses (I/I_Con_, +120 mV) for 8C (8D_NT_) variants to 1 mM ChlT (green symbols) and post-ox. DCPIB (orange symbols). I2Y bleach (1 mM) and post-bleach ChlT responses are included at right. Asterisks indicate statistical significance vs. pre-ChlT control values; crosses (+) indicate significance between post-ox. DCPIB and ChlT current level (***, P < 0.0005; **, ++ P < 0.005; * P < 0.05).

We then focused on the functional impact of residue I2 (8C) vs. F2 (8D) (Fig 1A). While both are hydrophobic, F2 is bulkier and also potentially able to interact with or modify the reactivity of adjacent M1 (Valley et al., 2012; Aledo et al., 2015; Chatterjee and Das, 2021; Chatterjee and Das, 2022). 8C I2F substitution resulted in current amplitudes similar to 8C WT (Fig. 5B, E), yet I2F currents displayed ChlT-dependent activation, although to a milder degree than 8C*_8D 2-4_*, exhibiting a 3.5-fold overall increase (P < 0.0005 vs. control; Fig. 5B). Finally, we replaced I2 with tyrosine (I2Y) to test whether the addition of a polar -OH would alter I2F behavior. I2Y-expressing cells had strongly reduced basal current amplitudes (1.90 ± 1.0 nA; n = 18) that displayed profound current activation in response to 1 mM ChlT, exhibiting a >50-fold overall increase over control levels (Fig. 5C). For all of the 8C-to-8D NT substitutions (8C_8D 1-22_, 8C_8D 2-4_, 8C_8D 5-11_, I2F, I2Y), the strength of ChlT-mediated current activation in individual recordings was similarly correlated with initial control current level (Fig. 6). The relationships for the five constructs distribute across a continuum, with I2F demonstrating larger currents and weakest activation, and I2Y displaying smallest currents and strongest relative activation (Fig. 6A, B and Supplemental Fig. S3).

**Figure 6.**
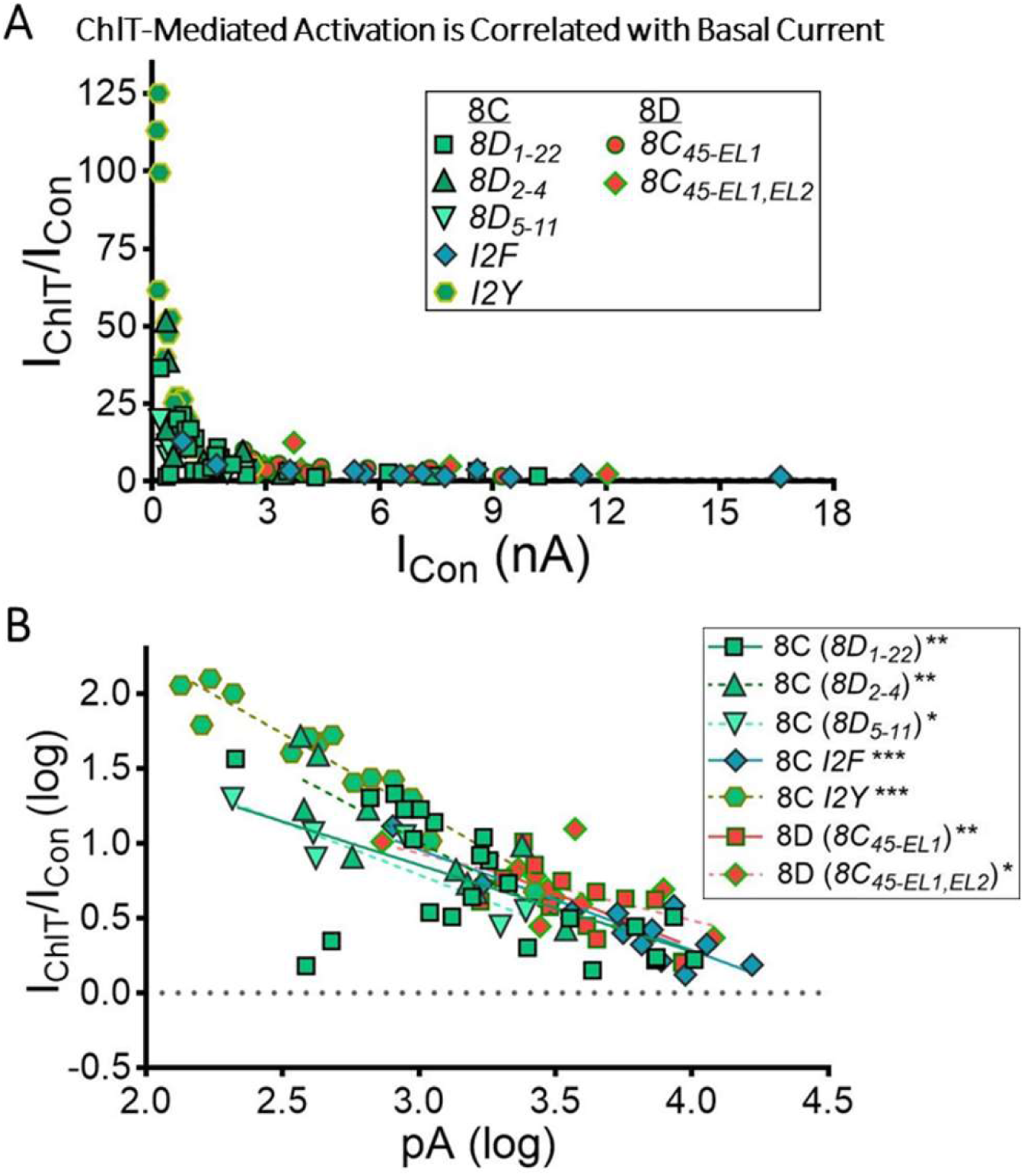
ChlT-mediated current activation is correlated with basal current amplitude. ***A,*** Relationship between current activation in response to 1 mM ChlT (I_ChlT_/I_Con_) and isotonic control current level (I_Con_, +120 mV), plotted for 8C and 8D constructs exhibiting activating responses. Each point represents an individual cell recording. ***B,*** Same data plotted on log scale (1.0 value corresponds to 10-fold activation). Lines fitted to each data group by linear regression demonstrate significant correlation with initial current (* P < 0.05; ** P < 0.005; P < 0.001).

Tyrosine residues can be chlorinated by ChlT or NaOCl (bleach)-derived HOCl (Hawkins et al., 2003; Woods et al., 2003; How et al., 2017; Hawkins, 2019; Nybo et al., 2019). To test the possibility that I2Y current activation is the result of Y2 chlorination, we recorded I2Y current responses to 5’ bleach application, followed by 1 mM ChlT application (Fig. 5D). Currents were unresponsive to 0.13 mM (94% of control; n = 3) or 1 mM bleach (92% of control; n = 3), but were activated by subsequent ChlT exposure (Fig. 5D). However, ChlT-mediated current activation following bleach treatment was weakened in a concentration-dependent manner (Fig. 5D,c and 5G), suggesting that bleach partially masked the activating action of ChlT.

In parallel, we assayed the current response of the 8D channel substituted with the 8C NT (8D*_8C 1-22_*). 8D*_8C 1-22_*-expressing cells demonstrated weak membrane expression, and small isotonic current amplitudes only 15% that of 8D WT (710 ± 45 pA; n = 8; Table 1). Currents in 5/6 cells displayed inhibitory responses to ChlT (33% to 72% inhibition vs. Iso control), characterized by a rapid maximal inhibition within 1’ ChlT (39 ± 9 % inhibition vs. control), but diminishing inhibition thereafter (29 ± 10% inhibition at 3’ ChlT). 8C NT substitution thus appeared to weaken 8D inhibitory sensitivity, but this assessment was complicated by the substantial reduction in initial current. This contrasts with 8D NT_2-11_ “out-of-context” substitutions in 8C that reverse the polarity of the WT 8C ChlT response, without modifying responses to other oxidants. This altered sensitivity appears to be regulated primarily by 2^nd^ NT residue.

### TM1 M48 regulates LRRC8C current inhibition and activation sensitivity

The TM1_23-44_ sequences of LRRC8A, 8C, and 8D are highly conserved, though 8C (C43) and 8D (M37) possess two differences in redox-active residues. However, distal TM1_45-49_ sequences differ in regard to methionine position (8A, LQVTQ; 8C, LQV**M**Q; 8D, **M**QLTK; Fig. 1A), with 8C M48 representing an oxidizable target at the apex of TM1 (Fig. 1C). As noted above, earlier LRRC8C domain analysis assigned the TM1_45-49_ sequence to the first extracellular loop (EL1). Notably, substitution of the entire 8C_45-EL1_ sequence to LRRC8A conferred robust VRAC activity to otherwise almost completely inactive 8A channels (Yamada and Strange, 2018b; Choi et al., 2021). We previously showed exchanging the 8C and 8D TM1_45-EL1_ sequences altered current responses to ChlT under hypotonic conditions, with 8C_45-EL1_ conferring activating responses to 8D, and 8D_45-EL1, EL2_ conferring 8D-like inhibitory sensitivity to 8C (Choi et al., 2021).

To clarify the contributions of TM1_45-49_ and EL domains in these current-modifying substitutions, we first reassessed this effect for 8C under isotonic conditions (Fig. 7A). 8D_45-EL1_ and 8D_45-EL1, EL2_ substitutions in 8C substantially reduced basal current level compared to WT (Fig. 7F and Table 1), presumably due to reduced channel trafficking to the cell surface. 8C_8D 45-EL1, EL2_-expressing cells had slightly stronger currents (2.5 ± 0.6 nA; n =8) which displayed rapid (30s-40s) and strongly inhibitory responses to ChlT, reminiscent of 8D WT (69 ± 4% inhibition vs. control; P < 0.001 vs. 8C WT; Fig. 7A, E). Substitution of 8D_45-49_ alone resulted in robust current amplitudes comparable to 8C WT (Fig. 7F), and enhanced ChlT-dependent inhibition (51 ± 4% inhibition vs. control, n = 7; P < 0.002 vs 8C WT; Fig. 7B, E). 8D_45-49_ substitution includes an M48T change, suggesting this switch supports increased inhibitory sensitivity. Consistent with this prediction, an M48T point substitution in 8C similarly enhanced ChlT-dependent current inhibition (53 ± 6% inhibition vs. control, n = 10; P < 0.05 vs 8C WT; Fig. 7C), although M48T current inhibition developed more rapidly than for 8C_8D 45-49_. (Fig. 7E).

**Figure 7.**
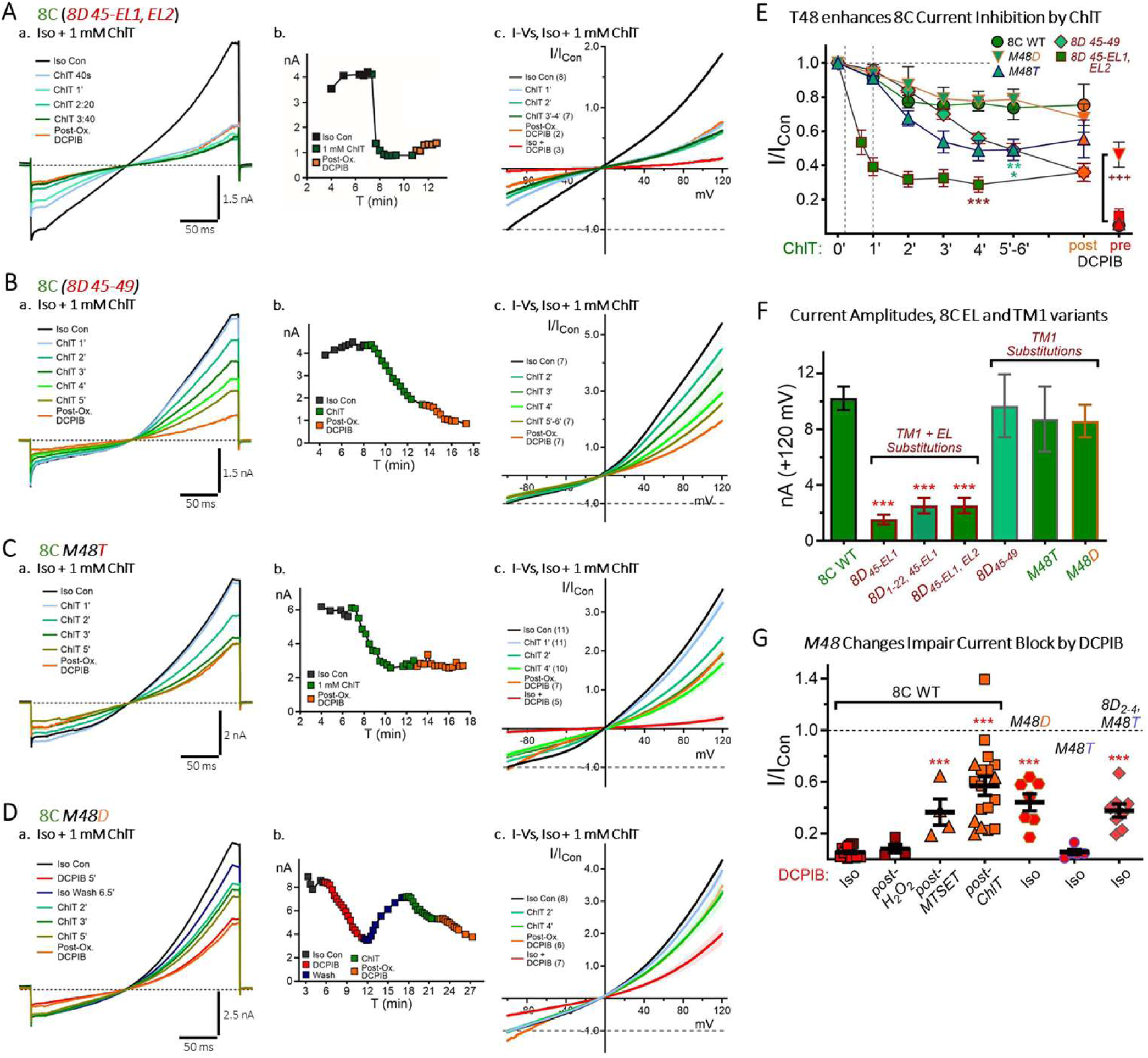
TM1_45-49_ and EL1 regulate LRRC8C inhibitory current response to ChlT. ***A-D***, ***Panel a***. Ramp-generated (−100 to +120 mV) isotonic currents during 1 mM ChlT application. Cells expressed GFP-tagged LRRC8C with the indicated 8D sequence substitution. ***Panel b***, Time course of current amplitude changes (+120 mV) during ChlT and 30 μM DCPIB application (orange, post-ChlT; red, pre-ChlT block) for the same cell. ***A,*** *8D45-el1,el2* (aa 45-163, 330-363) substitution results in rapid and potent current inhibition by ChlT, resembling the behavior of WT 8D currents (see Fig. 2B). ***B,*** *8D45-49* substitution results in enhanced (>50%) current inhibition compared to 8C WT. ***C*,** *M48T* substitution results in similarly enhanced (∼50%) inhibition compared to 8C WT. ***D***, *M48D* mutation displays incomplete current block by DCPIB (pre-ox. response, red trace/symbols). ChlT application results in moderate current inhibition similar to 8C WT. ***Panel c (A-D)***, Mean I-V relationships for each expression condition exposed to ChlT and DCPIB; currents are normalized to Iso control level at −100 mV (dotted lines). ***E***, Mean normalized current responses (I/I_Con_, +120 mV) to 1 mM ChlT (green symbols) and DCPIB (red, pre-ox; orange, post-ChlT) for the 8C constructs displayed in ***A-D***, compared to 8C WT. T48 conditions (*8D45-49* and *M48T*) significantly strengthened 8C inhibitory response at 4’-5’ (*** P < 0.0005; ** P < 0.005; * P < 0.05). Inclusion of the 8D EL1,2 domains was necessary to confer 8D-like ChlT response (*8D45-el1,el2*; squares). Post-ChlT application of DCPIB did not significantly inhibit current (orange symbols; P < 0.0005 vs. pre-ox DCPIB) in either M48 (WT) or T48 mutant contexts. ***F***, Isotonic basal current amplitudes (+120 mV) for 8C TM1 and EL variants (± S.E.M.). 8D EL substitutions (red outlines) resulted in significantly reduced basal current amplitude (***, P < 0.0005 vs 8C WT), limiting quantification of ChlT-mediated current inhibition to *8D45-el1,el2* (***A***). ***G***, Comparison of pre- and post-oxidant DCPIB current block for 8C WT and M48D and M48T mutants. Both *M48D* and 8D_2-4_, M48T conditions impaired DCPIB block to a degree comparable to that in 8C WT following ChlT and MTSET exposure (***, P < 0.0005 vs 8C WT pre-ox DCPIB).

These results suggested that overall inhibition for WT 8C is limited by ChlT-mediated oxidation of M48. We assessed an M48D mutation, which substitutes a negatively charged carboxyl that may mimick the effect of constitutive M48 oxidation to methionine sulfoxide (Fig. 7D). M48D currents were not significantly altered in basal amplitude or appearance (Fig. 7Da, 7F), and ChlT produced modestly inhibitory current responses comparable to those in 8C WT (21 ± 5% inhibition vs control, 4’ ChlT, n = 8; Fig. 7D-F). However, M48D current block by 30 mM DCPIB was significantly impaired compared to WT (n = 8; 56 ± 8% vs. 95 ± 1 % inhibition vs. 8C WT; Fig. 7Db,c and 7G). This result is consistent with the recent report that an 8C M48D mutation reduced binding and current inhibition by a lipophilic channel pore blocker, zafirlukast (Yamada et al., 2025). The level of M48D current inhibition by DCPIB did not differ significantly from WT post-ChlT DCPIB block (Fig. 7G), consistent with the idea that M48 oxidation impairs channel block. However, we observed that post-ChlT DCPIB block was also largely lost for 8D_45-49_ and M48T substitutions (Fig. 7B-C, E), indicating that impaired current block by DCPIB cannot be solely due to M48 oxidation, since it can also occur in theseT48 contexts.

Since M48T substitution enhanced 8C current inhibition by ChlT, we tested whether T48 also limits current activation resulting from 8D NT_2-4_ substitutions (Fig. 8). Indeed, the additional M48T substitution effectively eliminated current-activating responses in 8C_8D 2-4_, resulting in net current inhibition (12 ± 4% inhibition vs. Iso control; n = 8; Fig. 8A, a-b).

**Figure 8.**
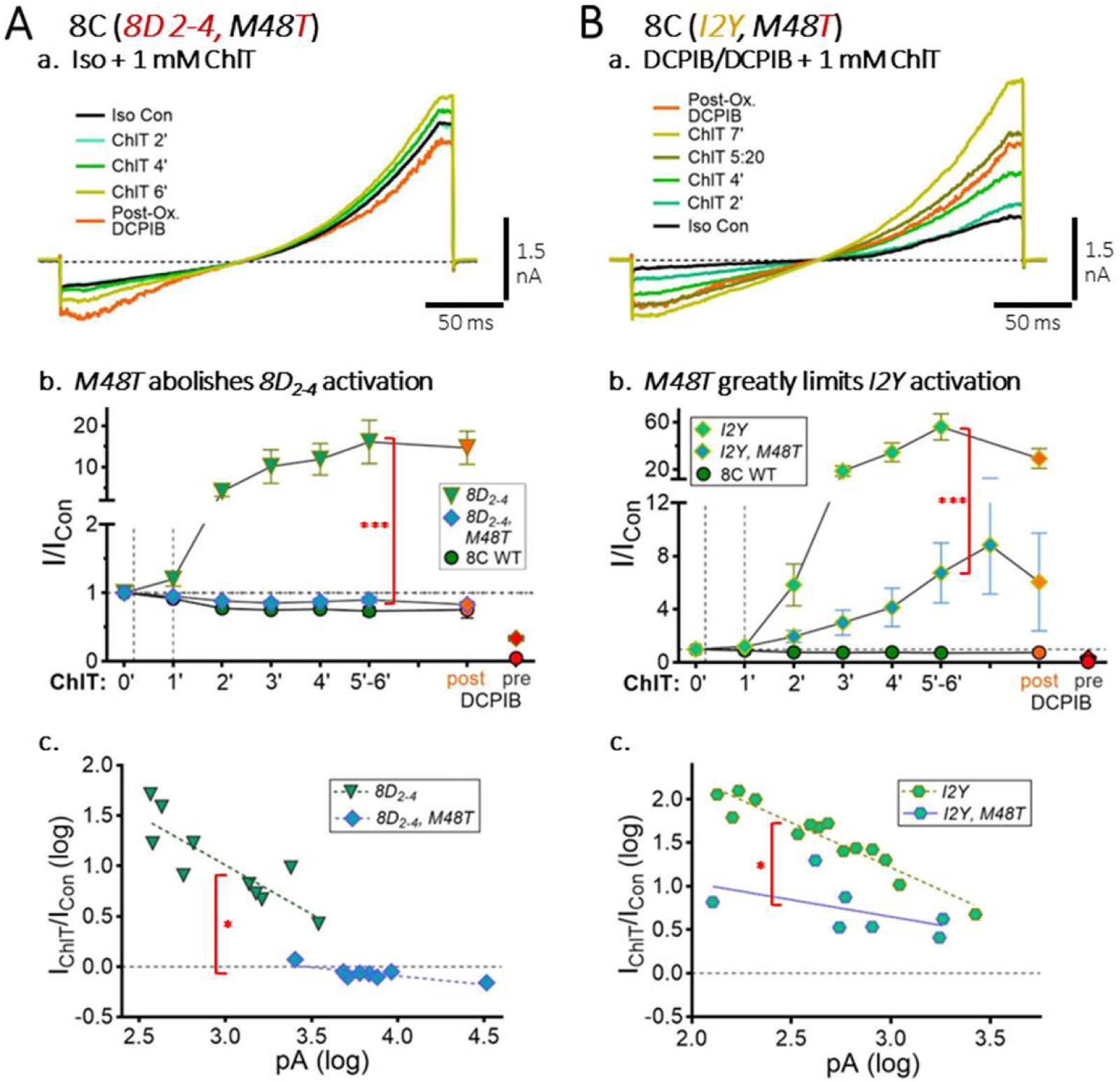
M48 is permissive for ChlT-mediated current activation in LRRC8C. The *M48T* substitution was added to 8C*_8D 2-4_* (***A***) and *I2Y* (***B***) backgrounds, both of which exhibit strong ChlT-dependent current activation. ***Panel a***, Ramp-generated (−100 to +120 mV) currents recorded under Iso control conditions (black trace) and after 1 mM ChlT addition (2’-8’). An activating response is nearly absent in 8C*_8D 2-4, M48T_* (***Aa***), while *I2Y, M48T* displays modest activation. ***Panel b***, Quantified normalized time-dependent ChlT responses (I/I_Con_) for *M48T* and background conditions, compared to 8C WT. In 8C*_8D 2-4, M48T_* (***Ab***), the activating response is completely lost and currents are weakly inhibited overall. ***Bb***, *M48T* greatly reduces the *I2Y* activating response (*** P < 0.0005 vs. M48 background). ***Panel c,*** *M48T* alters the relationship between the ChlT current response and initial current amplitude (log-transformed values). *8D_2-4_, M48T* (***Ac***) results in significantly increased basal current amplitude (P < 0.0005) and reduced slope (*P < 0.02) vs. *8D_2-4_,* while *I2Y, M48T* (***Bc***) results in significantly reduced slope (* P < 0.02 vs. in I2Y).

Notably, this change was accompanied by a ∼7-fold increase in basal current amplitudes and a loss of amplitude-dependence in the current response to ChlT (Fig. 8Ac). Unexpectedly, 8C_8D 2-4_, *M48T* currents also showed reduced current block by DCPIB (62 ± 5 % inhibition vs. control, n = 8; P < 0.001 vs 8C M48T and 8C WT; Fig. 7G), suggesting that the combined disruptions to NT_2-4_ and M48 may impact current block similarly to the M48D change. The M48T substitution likewise strongly reduced current activation in 8C *I2Y* by 88% (P < 0.0005 vs. I2Y, n = 7; Fig. 8Ba,b). In this case, *I2Y, M48T* basal current amplitude was not significantly altered, but the amplitude-dependence of the ChlT response was lost (Fig 8Bc). These results indicate that M48 is permissive, if not sufficient for current activation, suggesting that M48 oxidation leads to this form of current modulation.

### 8D activating currents require 8C TM1_45-49_ and EL1 sequence

Consistent with our previous findings, substitution of the 8C_45-EL1_ sequence in 8D resulted in pronounced current activation in response to 1 mM ChlT (4-fold increase over pre-ChlT control, 5’-6’ ChlT; n = 10), as well as increased basal current amplitude (P < 0.005 vs. 8D WT, n = 16; Fig. 9A, E-F and Table 1). Current activation was evident within 1’ and typically reached a maximum level after 4’-6’ ChlT (Fig 9Ab). 8D*_8C 45-EL1_* currents were strongly blocked by 30 μM DCPIB (n = 8; Fig 9Ab,c and Table 1). However, as in each previous context above, DCPIB block following ChlT exposure was largely absent (n = 6; Fig. 9Ab-c). 8D*_8C 45-EL1_* currents were unresponsive to 0.5 mM H_2_O_2_, but post-H_2_O_2_ ChlT application resulted in robust current activation (4.9-fold over pre-ChlT control; n = 4; Fig. 9B). DCPIB demonstrated substantial current block following H_2_O_2_ exposure (n = 2), but block was again greatly reduced or eliminated following subsequent ChlT exposure (panel b, Fig. 9A, B). The additional substitution of 8C EL2 (8D*_8C 45-EL1, EL2_*) resulted in comparable current-activating responses to ChlT (5.1-fold increase over pre-ChlT control; Fig. 9E and Table 1).

**Figure 9.**
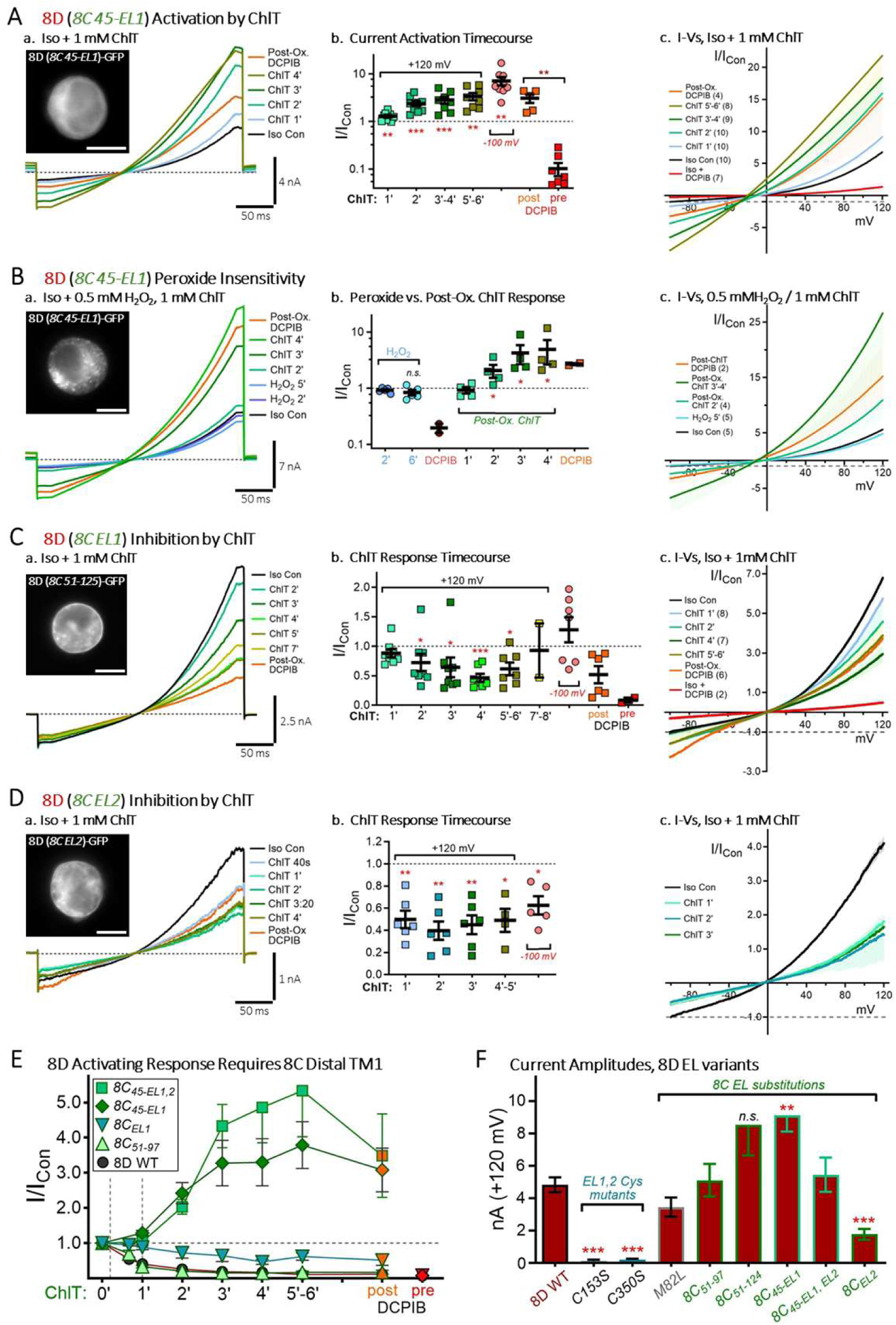
LRRC8C TM1_45-49_ + EL1 sequence confers current-activating ChlT responses to LRRC8D. ***A-D***, ***Panel a***. Ramp-generated (−100 to +120 mV) current responses to ChlT recorded from cells expressing GFP-tagged LRRC8D substituted with 8C*_45-EL1_* (aa 45-125; ***A-B***); 8C *EL1* (aa 51-125; ***C***); or 8C *EL2* (aa 286-317; ***D***). Images (***left***) illustrate protein distribution in the recorded cell (5 μm scale). ***Panel b***, Normalized current amplitude changes (I/I_Con_, +120 mV) during exposure to 1 mM ChlT (horizontal bracket) and 30 μM DCPIB (orange, post-ChlT; red, pre-ChlT block) in individual recordings; bars indicate mean values (± S.E.M.). Asterisks indicate significance vs. Iso or pre-ChlT control; crosses (+) indicate significance between pre-and post-ox. DCPIB (horizontal bracket; ***, P < 0.0005; **, ++ P < 0.005; * P < 0.05). ***A,*** 8D*_8C 45-EL1_* currents are activated several-fold over basal level by 3’-5’ ChlT application. The efficacy of DCPIB current block is greatly reduced or absent following ChlT exposure (orange vs. red symbols). ***B,*** 8D*_8C 45-EL1_* currents are insensitive to 0.5 mM H_2_O_2_ (5’; light blue trace/symbols), but strongly activated by subsequent ChlT exposure (green and tan traces/symbols). ***C*,** 8D*_8C EL1_* currents, by contrast, are markedly inhibited by ChlT for 3’-4’. Inhibition is diminished by longer exposure (6’-8’; tan and yellow traces/symbols). ***D***, 8D*_8C EL2_* expression resulted in smaller currents (see ***F***) that displayed overall moderately inhibitory responses to ChlT. ***Panel c (A-D)***, Mean I-V relationships for each expression condition exposed to ChlT (***A, C-D***), or H_2_O_2_ and ChlT (***B***). Current is normalized to Iso control level at - 100 mV (dotted line). ***E***, Time course of mean current responses (I/I_Con_, +120 mV; ± S.E.M.) to 1 mM ChlT (green symbols) and DCPIB (red, pre-ox; orange, post-ChlT) for 8D constructs substituted with 8C *EL1* including or excluding TM1_45-49_. 8C*_45-EL1_* and 8C*_45-EL1, EL2_* produced similar current-activating responses to ChlT. Substitution of 8C*_EL1_* or 8C*_EL2_* alone resulted in inhibitory responses. Post-ChlT current was not significantly inhibited by DCPIB. ***F***, Isotonic current amplitudes (+120 mV) for 8D with 8C_EL1,2_ + TM1 substitutions. 8D *C54S* (EL1) and *C353S* (EL2) cysteine mutations largely channel trafficking to the plasma membrane and had insignificant current. *M82L* exhibited essentially wild-type currents and inhibition, while 8C_51-97_ (partial EL1 substitution) retained strongly inhibitory current responses (see Table 1).

In contrast to the above results, substitution of 8C EL1 alone (8C_51-125_) in 8D resulted in inhibitory current responses to ChlT (8D*_8C EL1_*; Fig. 9C), but inhibition was less pronounced than for 8D WT (53 ± 8 % vs. control, 4’ ChlT, n = 8; P < 0.0005 vs 8D WT), and also decreased progressively at longer ChlT exposure times (5’-8’; Fig. 9Cb). Nevertheless, 8C EL1 substitution did not confer activating responses demonstrated by 8D*_8C 45-EL1_*. A partial 8C EL1 substitution (8C_51-97_) did not significantly alter 8D WT inhibitory behavior (Fig. 9E). 8C EL2 substitution alone resulted in weak current expression (1.56 ± 0.32 nA, P < 0.001 vs 8D WT; n = 9), but currents displayed inhibitory responses to ChlT, ruling out a current-activating action by EL2 (Fig. 9D). Overall, these results clarify that the distal portion of 8D EL1 (EL1_98-161_) contributes to the pronounced 8D inhibitory response to ChlT.

An M82L (EL1) mutant displayed WT current level and potent, WT-like current inhibition by ChlT (Fig. 9F and Table 1). C153S (EL1) and C350S (EL2) mutations respectively eliminate predicted C153-C335 and C54-C350 disulfide bonds linking EL1-EL2, which are required for proper protein trafficking. Predictably, these mutations eliminated significant currents in expressing cells (Fig. 9F and Table 1). EL1and EL2 substitutions, alone or in combination, resulted in increased current rectification, suggesting these domains contribute to the stronger current rectification displayed by WT 8C currents (Supplementary Fig. S2).

### TM1 T48M change disrupts 8D inhibitory current response to ChlT

The above results imply that the 8C TM1_45-49_ (LQV**M**Q) domain confers a current-activating ChlT response in the context of the 8D pore. We tested this prediction with the 8C_45-49_ substitution in 8D (Fig. 10A). Unexpectedly, 8D*_8C 45-49_* currents exhibited a multiphasic response to ChlT, characterized by rapid (1’) initial inhibition characteristic of 8D WT, followed by a progressive (1’-4’) increase of inward and outward currents (Fig. 10Ab), resulting in the loss of current inhibition within several minutes. Longer exposure was characterized by the further increase of poorly- or non-rectifying currents (Fig. 10Ac). Although current amplitudes at +120 mV were increased by several-fold after 8’-10’ ChlT exposure (Fig. 10Ae), the resultant “activated” currents differed markedly from ChlT-activated currents in the contexts described above (Figs. 4, 5, 9). 8D T48M substitution alone resulted in currents displaying stronger initial (1’-2’) current inhibition by ChlT and a more persistent overall inhibitory response (Fig. 10Ba,c,d). However, T48M currents exhibited the same progressive loss of rectification (Fig. 10Bb) and increased amplitude, leading to net current that exceeded initial control level after 6’-8’ (Fig. 10B, panel d). In contrast, 8D M45L currents displayed strong overall inhibition by ChlT (88 ± 5 % inhibition vs. Iso Con, 4’ ChlT; n = 5) that remained stable at 5’-6’ (Fig. 10C, E). Though the M45L current response had a slower inhibitory onset than 8D WT, the overall response suggests that M45 is not a major contributor to 8D inhibition. 8D TM1 has a charged Lys 49 residue at the apex of its helix, differing from Gln in 8A and 8C. We consequently tested an 8D K49Q substitution. Currents in K49Q expressing cells were strongly inhibited by ChlT (78 ± 13 % inhibition vs. Iso Con, 4’ ChlT; n = 3) (Fig. 10E and Table 1).

**Figure 10.**
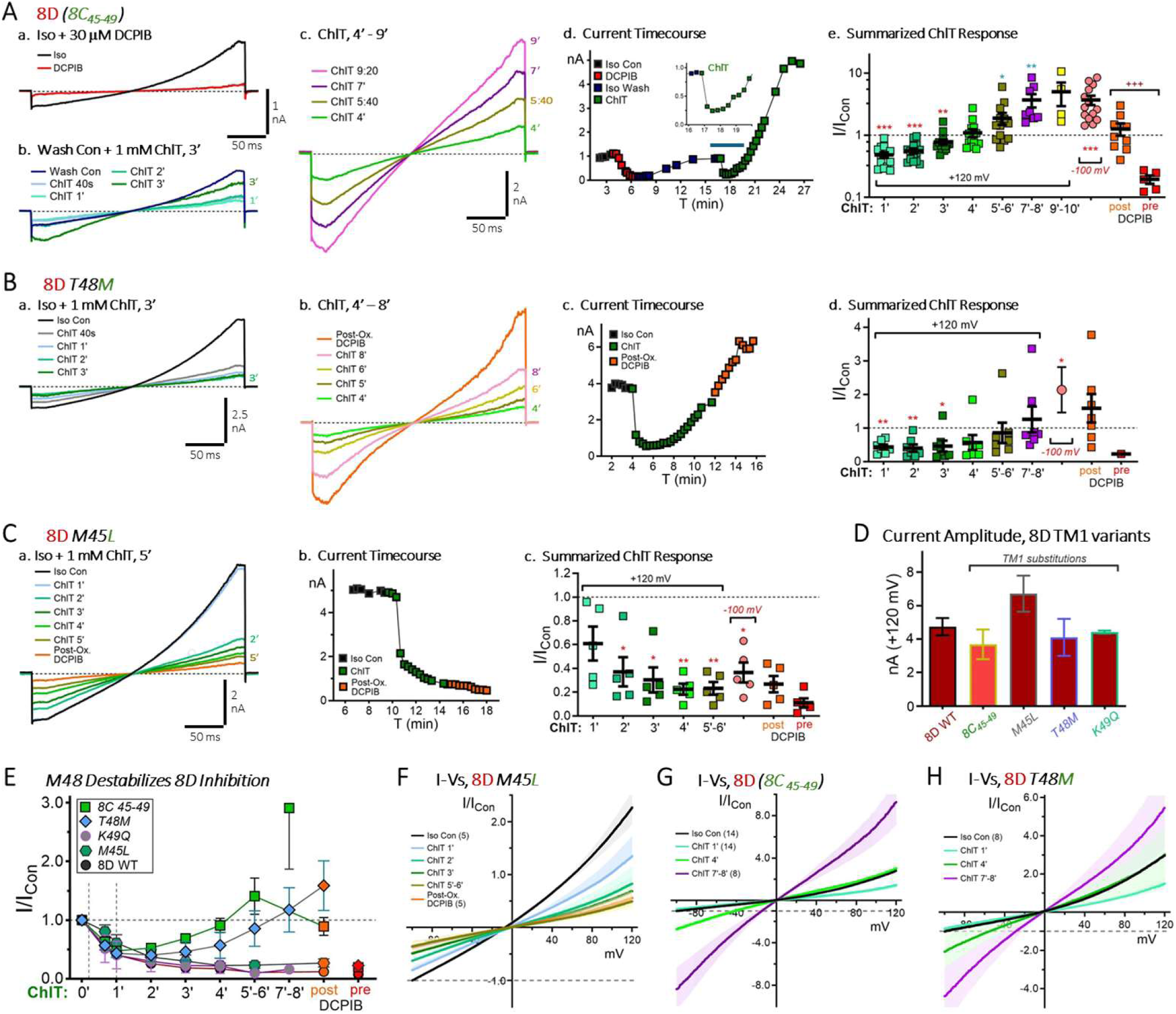
M48 destabilizes inhibitory ChlT action on LRRC8D current. ***A-C,*** Ramp currents (−100 to +120 mV) in cells expressed GFP-tagged LRRC8D substituted with the indicated 8C sequence. ***Aa,*** Isotonic 8D_8C *45-49*_ current is strongly blocked by 30 μM DCPIB. ***b,*** Following DCPIB washout, 1 mM ChlT application results in rapid (40-60s) initial current inhibition (ChlT 1’; light green trace); thereafter current amplitudes progressively increase (2’-3’ ChlT). ***c*,** Longer ChlT exposure (4’-9’) results in a progressive increase of non-rectifying current. ***Ad,*** Time course of current amplitude changes (+120 mV) for the cell shown in ***a-c***; inset shows the initial 3’ current response to ChlT in expanded time scale. ***Ae*,** Normalized current amplitude changes (I/I_Con_, +120 mV) for 8D_8C *45-49*_ during exposure to ChlT (horizontal bracket) and DCPIB (orange, post-ChlT; red, pre-ChlT block). Symbols: individual recordings; bars: mean values (± S.E.M). Mean current amplitude at 4’ ChlT exposure exceeded initial control level. Asterisks indicate significance vs. Iso Control; crosses (+) indicate significance between pre- and post-ox. DCPIB (***, +++ P < 0.0005; **, ++ P < 0.005; * P < 0.05). ***B,*** *T48M* current behaves similarly, displaying strong initial current inhibition by ChlT (1’-3’, ***a***), followed by the emergence and progressive increase of non-rectifying current (4’-8’ ChlT, ***b***). ***c,*** Time course of current amplitude changes (+120 mV) for the cell shown in ***a-b***. ***Bd*,** Normalized current amplitude changes (I/I_Con_, +120 mV) for *T48M.* Mean current amplitude exceeded initial control level after 6-8’ exposure to ChlT (significance indicated as in ***Ae****). **C,** M45L* current exhibits a pronounced and stable inhibitory response during 5’-6’ exposure to ChlT, resembling 8D WT. ***D***, Isotonic basal current amplitude (+120 mV) was not significantly altered by the TM1 substitutions assayed. ***E***, Time-dependence of current responses to ChlT for 8D TM1_45-49_ variants vs. 8D WT. M48 conditions (*8C_45-49_* and *T48M*) display a loss of initial current inhibition (40s - 2’), followed by an increasing atypical, non-rectifying current. Longer ChlT exposure (>4’-5’) resulted in net current increase. However, neither *8C_45-49_* or *T48M* supported current activation resembling 8D*_8C 45-EL1_* (see Fig. 9). ***F-H***, Mean normalized I-V relationships for the constructs illustrated in ***A-C***. 8C_45-49_ (***G***) and T48M (***H***) display initial inhibition (1’ ChlT; light green) followed by a subsequent loss of inhibition and increased inward current (4’ ChlT, dark green), and the progressive domination of a large non-rectifying current (7-8’ ChlT; magenta).

These results demonstrate that ChlT-mediated current inhibition in both 8C and 8D channels is highly sensitive to T48/M48. T48 substitution enhances 8C current inhibition, while M48 substitution in 8D destabilizes current inhibition and disrupts overall current features in response to ChlT. Nevertheless, both 8D EL1 and EL2 are required for its potent and stable inhibitory current response. Likewise, the conferral of recognizably activating current responses to 8D requires the presence of both 8C TM1_45-49_ as well as 8C EL1.

### ChlT disrupts DCPIB binding and channel block

Of various redox agents assayed, we find that only ChlT strongly modulates homomeric 8C and 8D channel currents. Additionally, ChlT action(s) disrupts current block by DCPIB in every post-ChlT context examined here and previously (Choi et al., 2021). Recent studies indicate that several LRRC8 channel blockers including DCPIB bind to lipid-associated domains within the channel pore, overlapping with TM1_45-49_ (Yamada et al., 2025). T48D/M48D mutations thought to result in pore delipidation (Kern et al., 2023) similarly impair current inhibition by blockers binding within the 8C channel pore (Yamada et al., 2025). These findings suggested that ChlT, while an effective oxidant and VRAC current modulator, may also occupy or otherwise interfere with this lipid-associated binding domain.

To support this hypothesis, we tested whether ChlT exposure disrupts pre-established DCPIB block of 8C and 8D channels. 30 μM DCPIB was applied in isotonic saline for several minutes to effect maximal (90%-95%) block of control current in three constructs (8C WT, 8C_8D 1-22_, and 8D_8C 45-EL1_) (Fig. 11A-C, panels a-b). 1 mM ChlT was then applied for 6’-10’ in the continued presence of DCPIB. In each context, current amplitudes progressively increased following ChlT addition, consistent with a degradation of channel block (panels a, c-d). The developing currents were poorly- or non-rectifying, differing from partially blocked currents of similar size during initial DCPIB application (Fig. 11A-C, panel d), or currents recovering from DCPIB block in the absence of ChlT (not shown). 8C_8D 1-22_ and 8D_8C 45-EL1_ developed larger currents in ChlT, suggesting some overall current activation over initial levels, even in the presence of DCPIB (Fig. 11D). WT 8C current is normally inhibited by ChlT, and current increase during ChlT exposure reached ∼50% of pre-block control level (Fig. 11D). Given the lipid/pore binding region proposed for other lipophilic blockers (Yamada et al., 2025), these results suggest that ChlT may disrupt the pore lipid environment to impair DCPIB binding efficacy.

**Figure 11.**
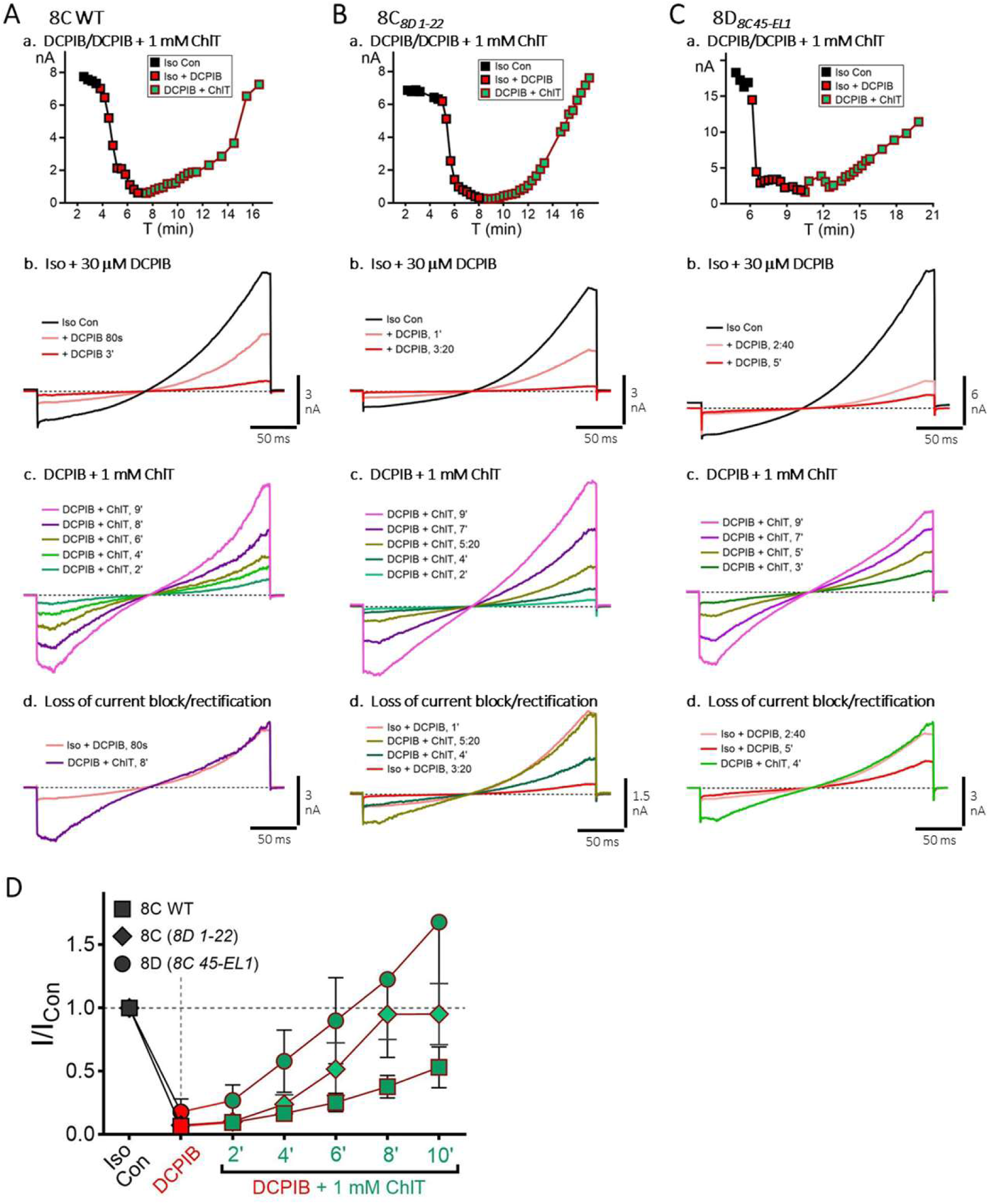
ChlT exposure disrupts and reverses existing LRRC8 channel block by DCPIB. Isotonic currents (−100 to +120 mV ramp) were recorded in cells expressing 8C WT (***A***), 8C*_8D 1-22_* (***B***), or 8D*_8C 45-EL1_* (***C***). DCPIB (30 μM) was applied for 2’-4’ to establish maximal current block (Iso + DCPIB; red symbols, pink/red traces). 1 mM ChlT was then added for 8’-10’ in the continued presence of DCPIB (DCPIB + ChlT; green symbols, green-pink traces). ***Panel a***, Time course of current amplitudes (+120 mV) recorded under control, DCPIB, and DCPIB + ChlT conditions. ***Panel b***, Current responses in Iso control saline and during DCPIB application. Pronounced current rectification is maintained during the onset of DCPIB block (pink traces). ***Panel c***, Current responses following addition of ChlT (same cells and scale as in ***a-b***). Current amplitudes increase progressively within 2’ exposure to ChlT (green traces), characterized by a loss of rectification (4’-9’). ***Panel d***, Comparison of current features during DCPIB block onset (pink, red) and following ChlT exposure (purple, tan, green), illustrating the appearance of inward currents and loss of rectification. ***D***, Timecourse of mean current amplitudes (I/ICon, +120 mV; ± S.E.M.) during initial DCPIB block (red symbols) and subsequent ChlT exposure (green symbols). Note that ChlT normally inhibits 8C WT current, but activates current for 8C*_8D 1-22_* and 8D*_8C 45-EL1_*. ChlT-dependent current increases for the latter two conditions may therefore reflect a combination of disrupted channel block and current activation.

## DISCUSSION

LRRC8D sequence differs notably from 8A/C/E family members within its proximal NT (NT_2-11_), distal TM1 helix (TM1_45-49_), and first extracellular loop (EL1_50-163_) domains. 8A/D and homomeric 8D channel currents are moreover distinguished by strongly inhibitory responses to ChlT (Gradogna et al., 2017b; Choi et al., 2021). Here we show that 8D NT substitution in 8C conferred strongly activating current responses to ChlT, reversing the WT inhibitory response. 8D_2-4_ substitutions, including the single I2F change, were sufficient to confer current activation to 8C, while I2Y resulted in maximally activating current responses to ChlT. In all contexts, relative activation was correlated to initial current amplitude, with smaller basal currents undergoing stronger fold-activation (Fig. 8). Basal current level likely reflects underlying channel activation level, in addition to other factors including channel trafficking and surface expression. Under isotonic conditions, 8C and 8D WT channels with C-terminal tags typically exhibit 30%-50% of maximal current activation produced by hypotonic saline (Gradogna et al., 2017b; Choi et al., 2021). Our results suggested a similar maximum capacity for ChlT-mediated current activation (20-25 nA). Importantly, 8C WT ChlT responses were inhibitory regardless of basal amplitude, while I2F currents invariably exhibited additional activation even when initial amplitude exceeded average WT level (Supplemental Fig. S3), indicating a fundamental shift in current modulation. Activating current responses required, or were greatly augmented by M48 in both native 8C and 8D (T48M) backgrounds. 8D current inhibition by ChlT was diminished by 8C EL1 substitution, supporting our previous conclusion (Choi et al., 2021) that the 8D inhibitory potency is partly conferred by its unique EL1. We conclude that NT_2_ has a fundamental role in regulating 8C channel modulation, while M48 oxidation limits inhibition and supports context-dependent activation.

### ChlT uniquely modulates VRAC current

ChlT is able to rapidly and significantly modulate 8C, and especially 8D currents that are partly activated by fluorescent tags or hypotonic saline (Gradogna et al., 2017b; Choi et al., 2021), and this study). Other oxidants (H_2_O_2_, TBHP, MTS reagents), NaOCl, and reducing agents (DTT, TCEP) exerted comparatively weak current inhibition or were without effect, and did not prevent subsequent responses to ChlT (Choi et al., 2021) and this study). ChlT thus possesses unique access or potency for VRAC current modulation. Only ChlT elicited current augmentation or activation under isotonic conditions, which has not been previously reported for 8C or 8D channels. TBHP and ChlT have been shown to activate 8A/E channel currents in oocytes and mammalian cells (Gradogna et al., 2017b; Bertelli et al., 2022), with the site of modulation mapped to intracellular LRD cysteines unique to LRRC8E (Bertelli et al., 2022).

ChlT effectively oxidizes protein M and C sulfurs (Shechter et al., 1975; Chandra Kumar et al., 1998; Dalle-Donne et al., 2002; Long et al., 2009; Yin et al., 2019), and can chlorinate various moieties via formation of HOCl, including the aromatic ring of tyrosine and phenylalanine, free amino groups, and lipid alkene groups (Ford, 2010; Schröter and Schiller, 2016; Tongul et al., 2018; Fricke et al., 2019; Knoke et al., 2024). The question remains whether ChlT or HOCl is able to access and oxidize M48 within the channel pore environment. Recent studies suggest that TM1 T48/M48 residues interact directly with the pore phospholipids that regulate 8A/C channel gating (Kern et al., 2023; Yamada et al., 2025). ChlT may therefore initiate oxidative N-chlorinating reactions with phospholipid head groups that subsequently oxidize associated protein residues, including M48 (Schröter and Schiller, 2016; Knoke et al., 2024). Though non-lipophilic, HOCl has been demonstrated to cause lipid peroxidation (Tongul et al., 2018). TBHP is lipophilic and also causes lipid peroxidation (Yang et al., 2020), while H_2_O_2_ is more polar than H_2_O and less directly toxic to membranes. Neither of these oxidants or NaOCl replicate the action of ChlT. It appears that ChlT possesses a relatively unique combination of properties that allow it to mediate these marked current-modulating effects.

### Role of the NT domain

8A/C channel function was shown to be strongly impaired by deletions and cysteine substitutions within the 8C NT_2-8_ region, as well as insertions which lengthen the NT while preserving NT_2-14_ sequence (Zhou et al., 2018). Channel function is thus highly sensitive to the precise positioning and interactions of NT_1-4_ within the pore. Our results suggest that 8D NT sequence substitutions disrupt 8C NT interactions, and/or displace NT_1-4_ from its folded position within the channel pore. It seems likely that the similar current-activating behavior produced by 8D NT_5-11_ substitution (TEFRQFS to AEVASLN) also results from a displacement of NT_1-4_. 8A hexameric channel structures have highly ordered NTs that are stabilized by E6-T5 and Y9-E8 polar interactions between dimer pairs, and non-polar interactions between I2-M37 and V4-I33/L36 of adjacent subunits, as well as NT-IL1,2 interactions that further stabilize pore structure (Liu et al., 2023). 8D NTs show less ordered structure, and do not form polar or non-polar interactions with neighboring NTs or TMs, consistent with greater NT flexibility and pore mobility (Nakamura et al., 2020; Liu et al., 2023). The 8C NT has not yet been clearly resolved in channel structures, but its predicted folded structure appears to closely resemble that of 8A (Yamada et al., 2025).

In the 8C pore environment, 8D NT_2-4_ substitutions (IPV to FTL) likely modify or eliminate native NT-NT and NT-TM1 interactions. It remains to be determined how such disruptions eliminate ChlT-dependent current inhibition and confer current activating responses. We speculate that this behavior is related to changes in NT stabilization via interactions with TM1 and TM2 residues. Compared to Isoleucine, aromatic groups form stronger interactions with an adjacent or neighboring Methionine via “methionine-aromatic interactions”, which yield additional stabilization energy (Valley et al., 2012; Yeung et al., 2020; Gibbs et al., 2021). M-oxidation eliminates the nonpolar I-M association, but oxidation strengthens methionine-aromatic interactions (Lewis et al., 2016; Walgenbach et al., 2018). These interactions can also alter redox reactivity of methionine in a 3D protein environment (Chatterjee and Das, 2021; Chatterjee and Das, 2022). Phenylalanine substitution (I2F) may further restrict NT by strengthening interaction with M37, and/or reduce M1 susceptibility to oxidation (Aledo et al., 2015) that normally favors current inhibition. Tyrosine (I2Y) forms an even stronger interaction with a greater impact on methionine reactivity. 8D_2-4_, 8D_5-11_, and I2Y substitutions resulted in smaller basal currents, suggesting that these NT changes decrease basal channel activation but confer a capacity for subsequent activation by ChlT. Notably, I2F and I2Y substitutions have remarkably different impacts on basal current level and relative activation. I2Y reduced basal current to nearly the level of deactivated channels in some cases (<300-400 pA), but ChlT exposure was able activate current to significantly above WT levels (Figs. 8, 12).

**Figure 12.**
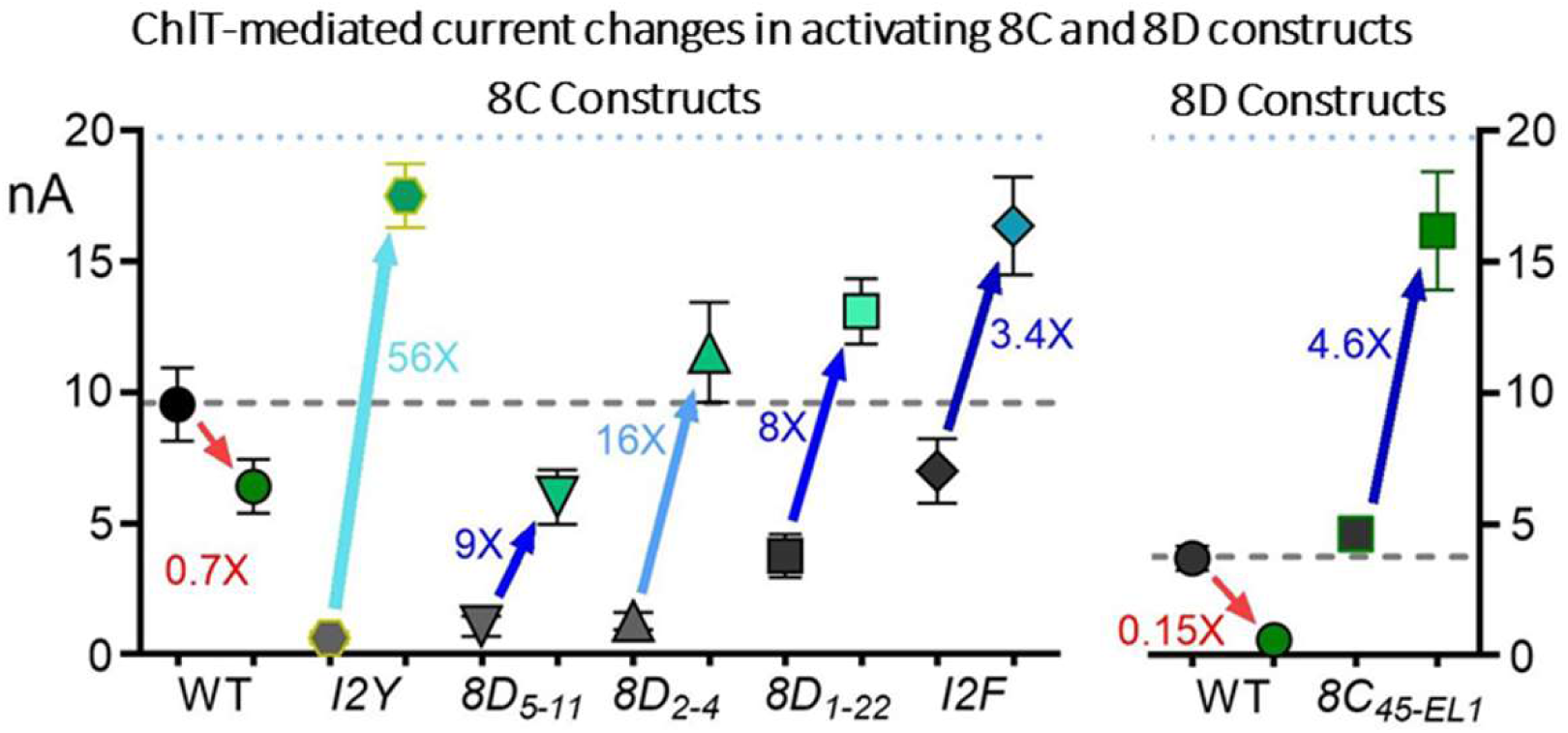
Mean basal and post-ChlT current levels for 8C (left plot) and 8D (right plot) channel contexts demonstrating activating responses to ChlT. Blue arrows/values indicate mean fold current activation; red arrows/values indicate current inhibition for WT 8C and 8D. Dashed grey lines represent WT basal current level,s and dotted blue lines approximate the maximal level of current activation suggested by recorded current values.

### Role of TM1, M48/T48, and EL1/2

The identity of TM1 48 (T48/M48) has a major impact on current sensitivity to ChlT. In 8D, activating current responses were conferred by the combined substitution of 8C TM1_45-49_ + 8C EL1, which included a T48M exchange as well as shortening of EL1 by 38 aa. Neither 8C_45-49_ or 8C EL1 individually conferred a similar effect. However, T48M alone resulted in the rapid disruption of 8D current inhibition, and the progressive “activation” of atypical non-rectifying currents, indicating that M48 is permissive and necessary for current activation. In 8C, M48T enhanced current inhibition by ChlT, and reversed or strongly reduced activating current responses resulting from 8D_NT 2-4_ or I2Y substitutions. These results clarify a functional role for T48 in stabilizing (8D) and/or enhancing (8C) inhibitory modulation. 8C M48T demonstrated enhanced (50%-60%) inhibition, comparable to that reported for 8A/C heteromeric channels (Gradogna et al., 2017b; Choi et al., 2021; Bertelli et al., 2022), which may reflect the presence of T48 in 8A subunits. M48 thus limits but does not prevent 8C current inhibition by ChlT, which has been attributed largely to M1 oxidation (Bertelli et al., 2022). However, when combined with NT_2-4_ disruption, for which I2F substitution is sufficient, M48 supports ChlT-dependent 8C current activation. Oxidation of M48 by ChlT is the most obvious explanation for the differences resulting from T48/M48 exchange.

ChlT exposure reduces or eliminates subsequent VRAC current block by DCPIB in every context examined, including WT and assorted mutant 8A, 8C, 8D, and 8E channel constructs and native VRAC currents (Choi et al., 2021); this study; and data not shown). M48D mutation reduced 8C DCPIB block efficacy to a similar degree (∼50%), consistent with the recent report that M48D impairs 8C current block by the lipophilic pore-binding agent zafirlukast (Yamada et al., 2025). However, the negatively charged M48D change may not be a good model for the sulfoxide expected to result from M48 oxidation. We also observed impaired post-ChlT current block by DCPIB in T48 contexts (8C_8D 45-49_ and M48T), ruling out M48 oxidation as the sole explanation for impaired block. A goal of future work will be to determine the identity of pore residue(s) that undergo oxidation or chlorination during exposure to ChlT.

The EL1 domain of 8D (123 aa) and 8C (75 aa) are 68% identical over their last 28 aa, but otherwise differ substantially (Choi et al., 2021). 8D EL1 possesses two regions (V75-T102, A121-N124) that absent in 8C and 8A, as well as a unique charge-rich (QEAKKEKK) sequence (aa 125-132). The 8D inhibitory response to ChlT was unaffected by substitution of 8C_51-97_, but diminished by over 50% by the complete exchange of 8C EL1, suggesting that the distal 66 aa of 8D EL1 contributes to its potent inhibition. This result supports our previous conclusion (Choi et al 2021) that 8D signature inhibition is partly conferred by its unique EL1.

### The lipidated channel microenvironment

Lipid densities aligned with the outer bilayer leaflet are closely associated with 8A/C and 8C channels structures (Kern et al., 2023; Takahashi et al., 2023). These densities were well-modeled by an external phospholipid packed into each subunit interface, with an acyl chain penetrating the inter-subunit cleft, and three internal phospholipids embedded in the channel pore, blocking ion flow in the closed channel configuration (Kern et al., 2023). The internal lipid polar head groups locate directly above T48/M48 residues at the apex of the TM1 helix, with their acyl chains lining the hydrophobic TM1 surfaces, extending two helical turns to approximately V40 (Kern et al., 2023; Yamada et al., 2025). Mutations in 8A introducing charged side groups to hydrophobic TM1 residues within TM1_40-48_ (eg, T48D, V47D/K, T44D, V40D) increased basal isotonic conductance, indicating reduced lipid occupancy and loss of closed-channel block (Kern et al., 2023; Yamada et al., 2025), while the T48D mutation resulted in a largely delipidated 8A/C channel that was associated with constitutively active current (Kern et al., 2023). Homomeric 8C-8A IL channels demonstrate heptameric structures with looser subunit arrangement than that of 8A/C channels (Takahashi et al., 2023; Yamada et al., 2025).

The lipophilic VRAC blocker zafirlukast was predicted to bind within the lipid-associated region of the 8C-8A IL pore, interacting with C43 and M48 of TM1, as well as hydrophobic TM2 residues (Yamada et al., 2025). It was suggested that zafirlukast binding may displace the flanking phospholipid acyl chains to induce current block. Modeling of predicted 8C NT structure within the pore suggested that the NT partially overlaps with the zafirlukast binding site (Yamada et al., 2025).

The C-terminal tagged channel constructs we have utilized are constitutively active to varying degrees (Choi et al., 2021), consistent with a partially expanded and conductive channel structure under isotonic conditions (Kern et al., 2023; Liu et al., 2023). The integrity of pore-associated lipids under isotonic conditions is supported by the effective current block demonstrated by DCPIB, which presumably is facilitated by its interaction with lipids or hydrophobic pore residues. The action of ChlT over a time course of minutes may induce disruption or retreat of pore lipids, leading to increased current and altered rectification properties. This behavior was most evident when M48 was introduced into the 8D NT/TM1 pore environment (T48M and 8C _45-49_ substitutions), resulting in rapid (2’-3’) disruption of initial inhibition and the progressive “activation” of atypical non-rectifying currents (Fig. 10), resembling the expected result of partial channel delipidation (Kern et al., 2023). ChlT action clearly reduces the favorability of subsequent DCPIB binding and current block, consistent with a disruption of the DCPIB binding region. ChlT moreover is capable of expelling DCPIB or otherwise reversing established channel block (Fig. 11). The resultant non-rectifying currents strongly resemble those activated by ChlT in 8D T48M mutants. Future work will increasingly focus on how the channel lipid microenvironment regulates LRRC8C and 8D function and modulation by oxidants. The gating of the 8D pore, which is substantially larger than 8C (Nakamura et al., 2020; Liu et al., 2023), seems likely to require more phospholipid molecules, which may affect the inherent stability as well as redox reactivity of the closed channel state.

### Conclusions

VRAC channels exist in an oxidant-rich local environment and are important modifiers of superoxide production, influx, and related inflammatory signaling. Under partially activated conditions, the 8D channel demonstrates markedly stronger ChlT-dependent current inhibition compared to the 8C channel. Current response is influenced by multiple structural elements, including the NT, particularly the second amino acid; M48, located at the apex of TM1 in 8C; and the distal portion of the 8D EL1. ChlT is a more effective and potent modifier of VRAC current than other related oxidants, although its chemical actions on specific 8C and 8D targets remain unclear. ChlT also disrupts channel inhibition by DCPIB, suggesting that it may also react with the lipid gate of the pore, which is the site of binding of multiple lipophilic channel blockers.

## Acknowledgments

The authors thank J. Denton, T. Yamada, and E. Karakas for helpful discussion and suggestions.

---

**Supplemental Figure S1.**
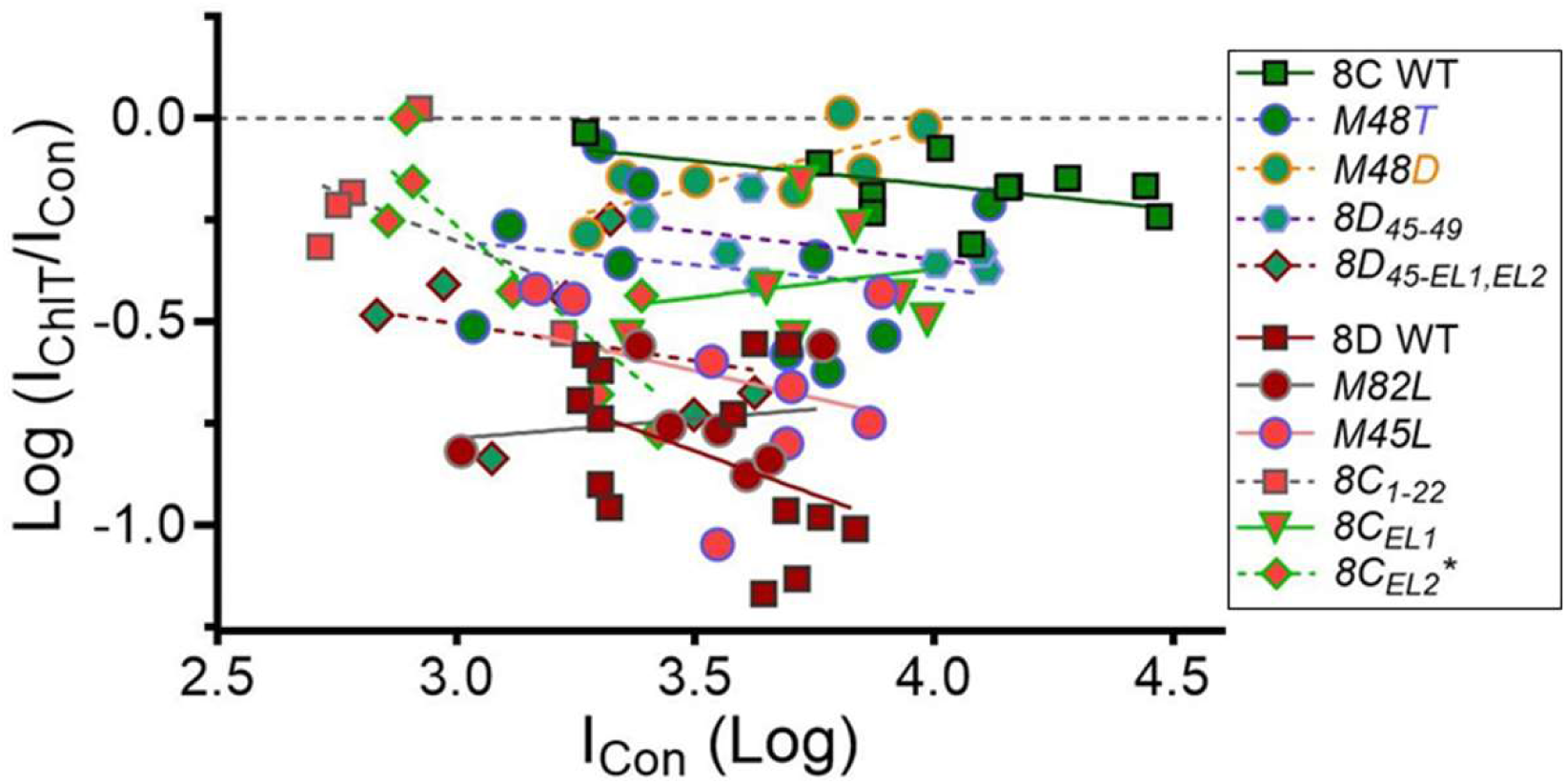
ChlT-mediated current inhibition is independent of basal current amplitude. Relative current change (I_ChlT_/I_Con_) in response to 1 mM ChlT is plotted against pre-ChlT isotonic control current (I_Con_, +120 mV) for LRRC8C and 8D constructs exhibiting inhibitory ChlT responses. Log value of −1.0 represents 90% current inhibition. Each point represents an individual cell recording. Lines were fit by linear regression to each data group. All slopes are insignificant, except for 8D_8C EL2_ which had small currents and displayed weak amplitude dependence (* P < 0.05; Dotted green line).

**Supplemental Figure S2.**
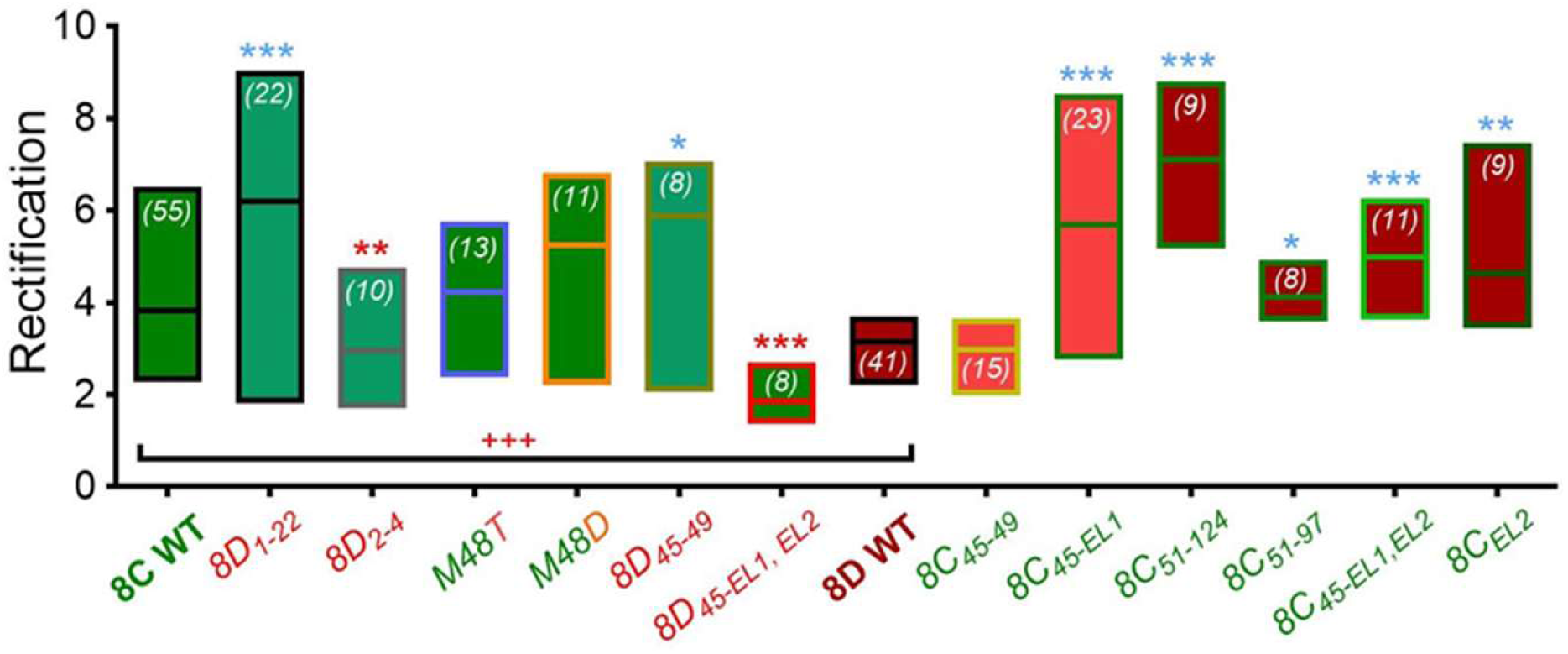
Current rectification changes resulting from NT, TM1, and EL substitutions. WT 8C ramp currents displayed stronger rectification (I_(+120 mV/_I_(−100 mV)_) than WT 8D (horizontal bracket; +++ P < 0.0005 vs. 8D). In the 8C background (green boxes), rectification was increased by substitution of the complete 8D NT (*8D_1-22_*) or TM1_45-49_ (*8D_45-49_*; blue asterisk), and decreased by 8D NT_2-4_ substitution (*8D_2-4_*; red asterisks). *8D_45-EL1, EL2_* substitution decreased rectification below the level of WT 8D. In the 8D background (red boxes), substitution of either or both 8C EL loops increased rectification (blue asterisks; * P < 0.05; ** P < 0.005; P < 0.001 vs. WT).

**Supplemental Figure S3.**
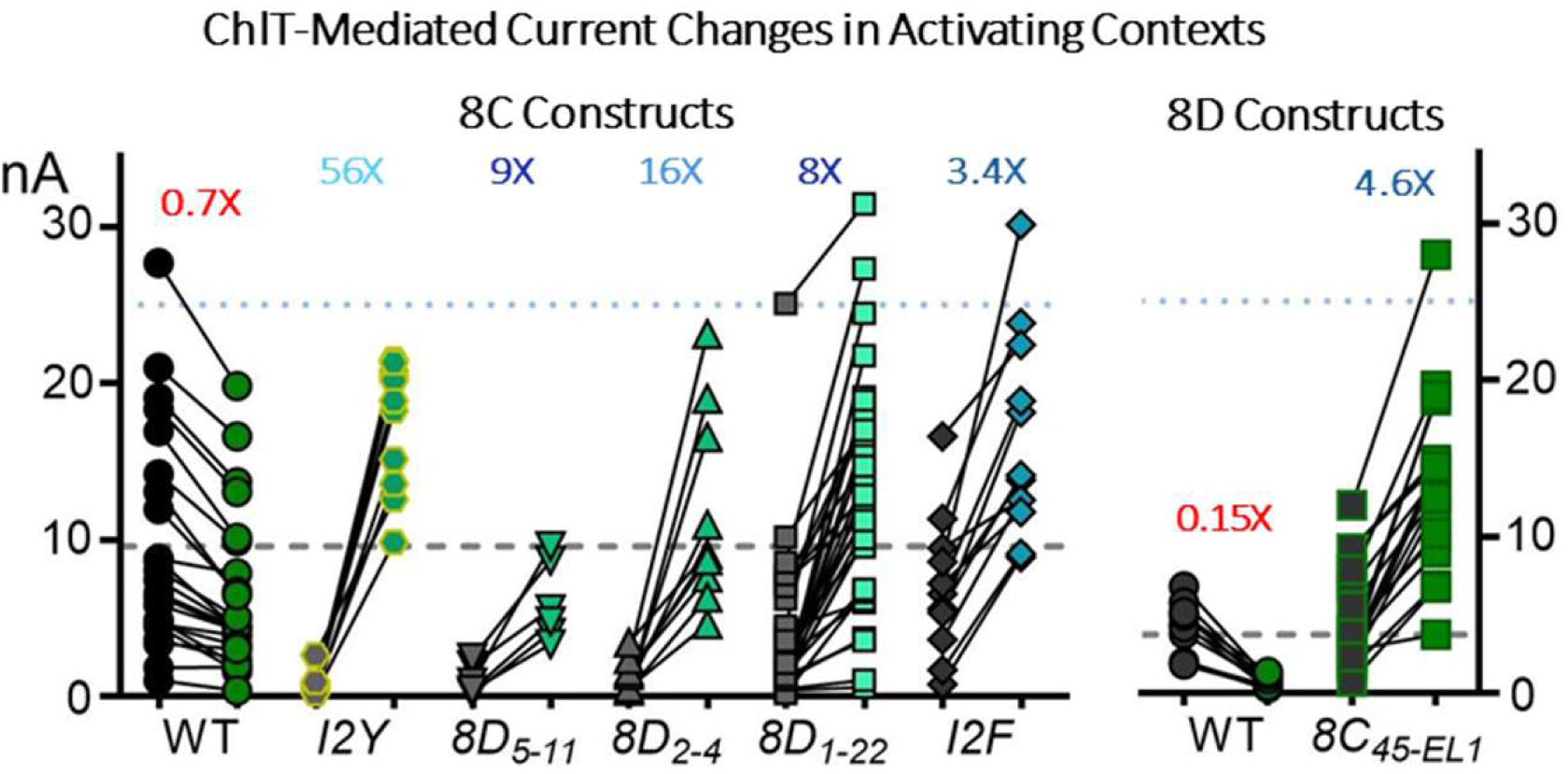
Basal control and post-ChlT current amplitudes (nA) recorded for 8C (left plot) and 8D channel constructs (right plot). Each pair of connected symbols represents an individual cell recording (left: control; right: 1 mM ChlT). Blue values indicate mean fold current activation (I_ChlT_/I_Con_). WT 8C and 8D currents displayed exclusively inhibitory responses (red values). Dashed grey lines represent mean WT basal current level. Dotted blue lines approximate the maximal level of current activation suggested by recorded values.

## References

1. Aledo, J.C., F.R. Cantón, and F.J. Veredas. 2015. Sulphur Atoms from Methionines Interacting with Aromatic Residues Are Less Prone to Oxidation. Sci Rep. 5:16955.

2. Bertelli, S., P. Zuccolini, P. Gavazzo, and M. Pusch. 2022. Molecular determinants underlying volume-regulated anion channel subunit-dependent oxidation sensitivity. J Physiol. 600:3965–3982.

3. Chandra Kumar, C., H. Nie, L. Armstrong, R. Zhang, S. Vijay-Kumar, and A. Tsarbopoulos. 1998. Chloramine T-induced structural and biochemical changes in echistatin. FEBS Letters. 429:239–248.

4. Chatterjee, K.S., and R. Das. 2021. An “up” oriented methionine-aromatic structural motif in SUMO is critical for its stability and activity. J Biol Chem. 297:100970.

5. Chatterjee, K.S., and R. Das. 2022. Methionine-aromatic Interaction as a Tool to Enhance Protein Stability. The FASEB Journal. 36.

6. Choi, H., N. Ettinger, J. Rohrbough, A. Dikalova, H.N. Nguyen, and F.S. Lamb. 2016. LRRC8A channels support TNFalpha-induced superoxide production by Nox1 which is required for receptor endocytosis. Free Radic Biol Med. 101:413–423.

7. Choi, H., M.R. Miller, H.N. Nguyen, J.C. Rohrbough, S.R. Koch, N. Boatwright, M.T. Yarboro, R. Sah, W.H. McDonald, J.J. Reese, R.J. Stark, and F.S. Lamb. 2023. LRRC8A anion channels modulate vascular reactivity via association with myosin phosphatase rho interacting protein. Faseb j. 37:e23028.

8. Choi, H., S. Panja, H.N. Nguyen, R.J. Stark, and F.S. Lamb. 2025. Smooth Muscle LRRC8A Knockout Preserves Vascular Function in Ang II Hypertension. Hypertension.

9. Choi, H., J.C. Rohrbough, H.N. Nguyen, A. Dikalova, and F.S. Lamb. 2021. Oxidant-resistant LRRC8A/C anion channels support superoxide production by NADPH oxidase 1. J Physiol. 599:3013–3036.

10. Dalle-Donne, I., R. Rossi, D. Giustarini, N. Gagliano, P. Di Simplicio, R. Colombo, and A. Milzani. 2002. Methionine oxidation as a major cause of the functional impairment of oxidized actin. Free Radical Biology and Medicine. 32:927–937.

11. Ford, D.A. 2010. Lipid oxidation by hypochlorous acid: chlorinated lipids in atherosclerosis and myocardial ischemia. Clin Lipidol. 5:835–852.

12. Friard, J., A. Laurain, I. Rubera, and C. Duranton. 2021. LRRC8/VRAC Channels and the Redox Balance: A Complex Relationship. Cell Physiol Biochem. 55:106–118.

13. Fricke, T.C., F. Echtermeyer, J. Zielke, J. de la Roche, M.R. Filipovic, S. Claverol, C. Herzog, M. Tominaga, R.A. Pumroy, V.Y. Moiseenkova-Bell, P.M. Zygmunt, A. Leffler, and M.J. Eberhardt. 2019. Oxidation of methionine residues activates the high-threshold heat-sensitive ion channel TRPV2. Proc Natl Acad Sci U S A. 116:24359–24365.

14. Gaitán-Peñas, H., A. Gradogna, L. Laparra-Cuervo, C. Solsona, V. Fernández-Dueñas, A. Barrallo-Gimeno, F. Ciruela, M. Lakadamyali, M. Pusch, and R. Estévez. 2016. Investigation of LRRC8-Mediated Volume-Regulated Anion Currents in Xenopus Oocytes. Biophys J. 111:1429–1443.

15. Gibbs, C.A., D.S. Weber, and J.J. Warren. 2021. Clustering of Aromatic Amino Acid Residues around Methionine in Proteins. Biomolecules. 12.

16. Gradogna, A., H. Gaitán-Peñas, A. Boccaccio, R. Estévez, and M. Pusch. 2017a. Cisplatin activates volume sensitive LRRC8 channel mediated currents in Xenopus oocytes. Channels (Austin). 11:254–260.

17. Gradogna, A., P. Gavazzo, A. Boccaccio, and M. Pusch. 2017b. Subunit-dependent oxidative stress sensitivity of LRRC8 volume-regulated anion channels. J Physiol. 595:6719–6733.

18. Harris, K., S.J. Won, G. Uruk, N. Mai, D. Ogut, Y. Zhao, L. Xie, P. Baxter, K. Everaerts, R. Sah, and R.A. Swanson. 2025. Excitotoxic neuronal death requires superoxide entry into neurons through volume-regulated anion channels. Sci Adv. 11:eadw0424.

19. Hawkins, Clare L. 2019. Hypochlorous acid-mediated modification of proteins and its consequences. Essays in Biochemistry. 64:75–86.

20. Hawkins, C.L., D.I. Pattison, and M.J. Davies. 2003. Hypochlorite-induced oxidation of amino acids, peptides and proteins. Amino Acids. 25:259–274.

21. How, Z.T., K.L. Linge, F. Busetti, and C.A. Joll. 2017. Chlorination of Amino Acids: Reaction Pathways and Reaction Rates. Environmental Science & Technology. 51:4870–4876.

22. Huo, C., Y. Liu, X. Li, R. Xu, X. Jia, L. Hou, and X. Wang. 2021. LRRC8A contributes to angiotensin II-induced cardiac hypertrophy by interacting with NADPH oxidases via the C-terminal leucine-rich repeat domain. Free Radic Biol Med. 165:191–202.

23. Kern, D.M., J. Bleier, S. Mukherjee, J.M. Hill, A.A. Kossiakoff, E.Y. Isacoff, and S.G. Brohawn. 2023. Structural basis for assembly and lipid-mediated gating of LRRC8A:C volume-regulated anion channels. Nat Struct Mol Biol. 30:841–852.

24. Knoke, L.R., S.A. Herrera, S. Heinrich, F.M.L. Peeters, N. Lupilov, J.E. Bandow, and T.G. Pomorski. 2024. HOCl forms lipid N-chloramines in cell membranes of bacteria and immune cells. Free Radic Biol Med. 224:588–599.

25. Lamb, F.S., H. Choi, M.R. Miller, and R.J. Stark. 2024. Vascular Inflammation and Smooth Muscle Contractility: The Role of Nox1-Derived Superoxide and LRRC8 Anion Channels. Hypertension.

26. Lewis, A.K., K.M. Dunleavy, T.L. Senkow, C. Her, B.T. Horn, M.A. Jersett, R. Mahling, M.R. McCarthy, G.T. Perell, C.C. Valley, C.B. Karim, J. Gao, W.C.K. Pomerantz, D.D. Thomas, A. Cembran, A. Hinderliter, and J.N. Sachs. 2016. Oxidation increases the strength of the methionine-aromatic interaction. Nature Chemical Biology. 12:860–866.

27. Liu, H., M.M. Polovitskaya, L. Yang, M. Li, H. Li, Z. Han, J. Wu, Q. Zhang, T.J. Jentsch, and J. Liao. 2023. Structural insights into anion selectivity and activation mechanism of LRRC8 volume-regulated anion channels. Cell Rep. 42:112926.

28. Long, L.H., J. Liu, R.L. Liu, F. Wang, Z.L. Hu, N. Xie, H. Fu, and J.G. Chen. 2009. Differential effects of methionine and cysteine oxidation on [Ca2+] i in cultured hippocampal neurons. Cell Mol Neurobiol. 29:7–15.

29. Matsuda, J.J., M.S. Filali, J.G. Moreland, F.J. Miller, and F.S. Lamb. 2010. Activation of swelling-activated chloride current by tumor necrosis factor-alpha requires ClC-3-dependent endosomal reactive oxygen production. J Biol Chem. 285:22864–22873.

30. Nakamura, R., T. Numata, G. Kasuya, T. Yokoyama, T. Nishizawa, T. Kusakizako, T. Kato, T. Hagino, N. Dohmae, M. Inoue, K. Watanabe, H. Ichijo, M. Kikkawa, M. Shirouzu, T.J. Jentsch, R. Ishitani, Y. Okada, and O. Nureki. 2020. Cryo-EM structure of the volume-regulated anion channel LRRC8D isoform identifies features important for substrate permeation. Commun Biol. 3:240.

31. Nybo, T., M.J. Davies, and A. Rogowska-Wrzesinska. 2019. Analysis of protein chlorination by mass spectrometry. Redox Biology. 26:101236.

32. Panja, S., H. Choi, H.N. Nguyen, J. Russolillo, S. Dikalov, R.J. Stark, and F.S. Lamb. 2025. Smooth muscle LRRC8A knockout reduces O_2_^·-^ influx, inflammation, senescence and atherosclerosis. bioRxiv.2025.2012.2001.691680.

33. Quinodoz, M., S. Rutz, V. Peter, L. Garavelli, A.M. Innes, E.F. Lehmann, S. Kellenberger, Z. Peng, A. Barone, B. Campos-Xavier, S. Unger, C. Rivolta, R. Dutzler, and A. Superti-Furga. 2025. De novo variants in LRRC8C resulting in constitutive channel activation cause a human multisystem disorder. Embo j. 44:413–436.

34. Ren, Z., F.J. Raucci, Jr., D.M. Browe, and C.M. Baumgarten. 2008. Regulation of swelling-activated Cl(-) current by angiotensin II signalling and NADPH oxidase in rabbit ventricle. Cardiovasc Res. 77:73–80.

35. Rutz, S., D. Deneka, A. Dittmann, M. Sawicka, and R. Dutzler. 2023. Structure of a volume-regulated heteromeric LRRC8A/C channel. Nat Struct Mol Biol. 30:52–61.

36. Schröter, J., and J. Schiller. 2016. Chlorinated Phospholipids and Fatty Acids: (Patho)physiological Relevance, Potential Toxicity, and Analysis of Lipid Chlorohydrins. Oxid Med Cell Longev. 2016:8386362.

37. Shechter, Y., Y. Burstein, and A. Patchornik. 1975. Selective oxidation of methionine residues in proteins. Biochemistry. 14:4497–4503.

38. Takahashi, H., T. Yamada, J.S. Denton, K. Strange, and E. Karakas. 2023. Cryo-EM structures of an LRRC8 chimera with native functional properties reveal heptameric assembly. Elife. 12.

39. Thöne, F.M.B., M.M. Polovitskaya, and T.J. Jentsch. 2025. LRRC8/VRAC chloride and metabolite channels in signaling and volume regulation. Trends Biochem Sci. 50:873–891.

40. Tongul, B., B. Kavakcıoğlu, and L. Tarhan. 2018. Chloramine T induced oxidative stress and the response of antioxidant system in Phanerochaete chrysosporium. Folia Microbiol (Praha). 63:325–333.

41. Valley, C.C., A. Cembran, J.D. Perlmutter, A.K. Lewis, N.P. Labello, J. Gao, and J.N. Sachs. 2012. The methionine-aromatic motif plays a unique role in stabilizing protein structure. J Biol Chem. 287:34979–34991.

42. Varela, D., F. Simon, A. Riveros, F. Jørgensen, and A. Stutzin. 2004. NAD(P)H oxidase-derived H(2)O(2) signals chloride channel activation in cell volume regulation and cell proliferation. J Biol Chem. 279:13301–13304.

43. Walgenbach, D.G., A.J. Gregory, and J.C. Klein. 2018. Unique methionine-aromatic interactions govern the calmodulin redox sensor. Biochemical and Biophysical Research Communications. 505:236–241.

44. Woods, A.A., S.M. Linton, and M.J. Davies. 2003. Detection of HOCl-mediated protein oxidation products in the extracellular matrix of human atherosclerotic plaques. Biochemical Journal. 370:729–735.

45. Yamada, T., P. Bisignano, E. Karakas, and J.S. Denton. 2025. A conserved mechanism of LRRC8 channel inhibition by two structurally distinct drugs. Commun Biol. 8:1432.

46. Yamada, T., and K. Strange. 2018a. Intracellular and extracellular loops of LRRC8 are essential for volume-regulated anion channel function. J Gen Physiol.

47. Yamada, T., and K. Strange. 2018b. Intracellular and extracellular loops of LRRC8 are essential for volume-regulated anion channel function. J Gen Physiol. 150:1003–1015.

48. Yang, H.C., H. Yu, T.H. Ma, W.Y. Tjong, A. Stern, and D.T. Chiu. 2020. tert-Butyl Hydroperoxide (tBHP)-Induced Lipid Peroxidation and Embryonic Defects Resemble Glucose-6-Phosphate Dehydrogenase (G6PD) Deficiency in C. elegans. Int J Mol Sci. 21.

49. Yeung, P.S.W., C.E. Ing, M. Yamashita, R. Pomès, and M. Prakriya. 2020. A sulfur-aromatic gate latch is essential for opening of the Orai1 channel pore. eLife. 9:e60751.

50. Yin, V., S.H. Mian, and L. Konermann. 2019. Lysine carbonylation is a previously unrecognized contributor to peroxidase activation of cytochrome c by chloramine-T. Chem Sci. 10:2349–2359.

51. Zhou, P., M.M. Polovitskaya, and T.J. Jentsch. 2018. LRRC8 N termini influence pore properties and gating of volume-regulated anion channels (VRACs). J Biol Chem. 293:13440–13451.

